# Basal forebrain projections to the lateral habenula regulate ethanol and sucrose consumption

**DOI:** 10.64898/2026.07.02.736151

**Authors:** Klaiten Kermoade, Emaleigh Hulet, Anika Paulson, Porshia Woods, Gelila Woldemariam, Jocelyn M. Richard

## Abstract

**Background:** The lateral habenula (LHb) receives afferent input from various basal forebrain (BF) regions known for their role in aversion-related behaviors, depression, and addiction. However, little is known about the role of this circuitry in ethanol consumption, including in the presence of aversive stimuli. Here, we assessed the role of BF input to the LHb in alcohol and sucrose consumption using a chemogenetic approach.

**Methods:** Following long-term intermittent access to alcohol (IAA) or control conditions, male and female Long-Evans rats underwent surgery for the expression of designer receptors in BF-to-LHb projections. We then excited this pathway in rats with IAA history, or inhibited this pathway in rats with limited ethanol history, before testing consumption of ethanol and natural reward both in the presence and absence of quinine adulteration.

**Results:** Long-term IAA elevated ethanol drinking in all rats, and enhanced ethanol preference with and without quinine adulteration in males. Chemogenetic activation of BF-to-LHb neurons in rats with IAA history produced different effects in males and females: excitation enhanced ethanol intake in females, but reduced ethanol preference in males, regardless of quinine adulteration. Activation also led to a relative insensitivity to quinine adulteration of sucrose when compared to controls, particularly in females. Chemogenetic inhibition in rats with limited ethanol history did not alter consumption of ethanol or sucrose, with or without quinine adulteration.

**Conclusions:** Our results suggest a role for BF-to-LHb circuitry in ethanol drinking that may vary based on sex, and a potential role for this circuitry in the sensitivity to quinine when combined with natural reward.

## INTRODUCTION

The lateral habenula (LHb) is a well-conserved epithalamic nucleus responsible for encoding a wide variety of aversive stimuli (Groos & Helmchen, 2024; Hu et al., 2020) and constraint of reward seeking (Post et al., 2022). The LHb receives diverse afferent projections (Groos & Helmchen, 2024; Hikosaka et al., 2008). One important input is the basal forebrain (BF), a heterogeneous subcortical region implicated in various behaviors not limited to arousal, learning, and reward-related behavior (Heimer et al., 1993; Napier et al., 1991; Rieck et al., 1995), comprising multiple regions including the ventral pallidum (VP), diagonal band of Broca (DBB), and the lateral preoptic area (LPO), among others. Neurons projecting from the BF to LHb (BF-to-LHb) have been shown to signal noxious stimuli (Barker et al., 2017), drive place avoidance (Faget et al., 2018; Swanson et al., 2022), and regulate depression phenotypes (Knowland et al., 2017), while modulating both intake and seeking of addictive drugs, such as cocaine (Heinsbroek et al., 2020; Levi et al., 2025) and heroin (Chen et al., 2024). This projection is also well-positioned to mediate the effects of the nucleus accumbens (NAc) on aversion-resistant drinking, as BF subregions (in particular, but not limited to, the VP) comprise a major NAc output locus (Smith et al., 2009; Zhang et al., 2013; Soares-Cunha & Heinsbroek, 2023; Pan et al., 2024). Nevertheless, the role of BF-to-LHb projections in consumption of ethanol or sugar reward, in the presence or absence of aversive stimuli, remains unclear.

Here, we tested the effects of chemogenetic manipulation of BF-to-LHb projections in male and female Long-Evans rats, which we hypothesized would bidirectionally modulate ethanol consumption. We tested the role of this pathway in rats with two distinct alcohol exposure histories. One set of animals underwent a long-term 2-bottle choice (2-BC) intermittent 15% alcohol access (IAA) protocol to establish regular ethanol intake. This procedure has been used previously in our lab and others to establish motivation for alcohol prior to cue learning (Carpio et al., 2022; Cofresí et al., 2017, 2018, 2019; LeCocq et al., 2022; Millan et al., 2015, 2017; Segal et al., 2022; Sood et al., 2026; Sparks et al., 2014) and has been shown to promote ethanol reward seeking and reinforcement (Angelyn et al., 2021; Rodd et al., 2005). We also expected the emergence of aversion-resistant ethanol drinking following this paradigm, which has been found after using a similar procedure with 20% ethanol (Hopf et al., 2010; Seif et al., 2013, 2015; Simms et al., 2008; Wise, 1973). A separate group of animals received only minimal exposure to ethanol prior to testing. In these groups, we assessed whether we could alter ethanol or natural reward drinking through manipulation of the BF-to-LHb circuit; given the established role of BF-to-LHb projections in aversion, we also tested whether modulation of this circuit could alter intake in the presence of aversive stimuli (i.e., quinine adulteration). We hypothesized that chemogenetic activation of BF-to-LHb projections could suppress ethanol or sugar reward intake either with or without negative consequences in animals with protracted ethanol history. In contrast, we hypothesized that chemogenetic inhibition of BF-to-LHb projections could enhance ethanol or sugar reward consumption either with or without negative consequences in animals with limited ethanol history. Here, we found that activation of BF-to-LHb circuitry altered ethanol drinking behavior in Long-Evans rats following long-term IAA, with effects varying by sex, while diminishing quinine-driven suppression of sucrose intake.

## METHODS

### Subjects

Adult female and male Long-Evans rats were obtained from Envigo or an in-house colony and housed in a temperature-and humidity-controlled colony room. Females weighed ∼200-250 (∼P65-98) and males weighed ∼250-425 (∼P65-P99) upon arrival or the start of experiments. Rats obtained from Envigo were used for the primary experiments assessing the impact of long-term IAA and chemogenetic manipulations. Upon arrival these rats were initially pair-housed for 1 week on a 14:10 light/dark cycle, and then individually housed for the remainder of the experiment. Colony rats were used for chemogenetic ligand control experiments and were individually housed on a 12:12 light/dark cycle at the start of the experiment. Water and food was available *ad libitum* throughout the experiments. All experimental procedures were approved by the Institutional Animal Care and Use Committee at the University of Minnesota and were conducted in accordance with the guidelines on animal care and use of the National Institutes of Health of the United States.

### Long-term access to alcohol

Long-term access (LTA) rats (n=50, 23F/27M) received 2-bottle choice intermittent alcohol access (2-BC IAA) with 15% ethanol, comprising three 24-hour periods per week interleaved with 24-48 hour withdrawal periods to promote ethanol drinking. Water was available at all times. At week 12, rats received a 7-to-10-day abstinence period to complete intracranial viral surgery. Following surgery, rats were re-exposed to intermittent ethanol to reach a total of 14 weeks (42 sessions) of intermittent access. For direct comparison to LTA rats, a group of age-matched short-term access (STA) rats (n=24, 11F/13M) underwent identical procedures during this period except that they received a second water bottle in place of ethanol. Additional STA rats (n=34; 15F/19M) that were not age matched were also used to assess the impact of BF-to-LHb manipulations, but were never directly compared to LTA rats.

### Intracranial surgery

Rats were induced for anesthesia with isoflurane (5%) before transfer to a stereotaxic apparatus (David Kopf Instruments), where anesthesia was maintained at 0.5-2.5% (SomnoSuite). An intersectional dual-virus strategy was used to bilaterally target either chemogenetic receptors or control fluorophores to neurons projecting from the BF to LHb. Syringes for viral delivery were aimed between the following coordinates relative to bregma, from the skull surface — BF: +0.3 mm anteroposterior (A/P), ±2.30 - 2.50 mm mediolateral (M/L), -8.00 - 8.40 mm dorsoventral (D/V); LHb: -3.15 - 3.70 mm A/P, ±0.73 -0.83 mm M/L, - 5.05 - 5.90 mm D/V. LTA rats received infusions of both a retrograde AAV encoding Cre recombinase (Addgene 105553-AAVrg, pENN.AAV.hSyn.Cre.WPRE.hGH, 1.9 x10^13^ GC/mL, or AAV2retro-hSyn-Cre, prepared by University of Minnesota Viral Core, 3.0 x10^13^ GC/mL) into the LHb and Cre-dependent viruses expressing either designer Gq receptors (n = 18 (9F/9M); Addgene 44361-AAV8, pAAV-hSyn-DIO-hM3D(Gq)-mCherry, 2.1 x10^13^ GC/mL) or mCherry control (n = 14 (7F/7M); Addgene 50459-AAV8, pAAV-hSyn-DIO-mCherry, 1.7 - 2.2 x10^13^ GC/mL) into the BF. STA rats received infusions of both a Cre-expressing retrograde AAV into the LHb (listed above), and Cre-dependent viruses expressing either designer Gi receptors (n = 34, 15F/19M; Addgene 44362-AAV8, pAAV-hSyn-DIO-hM4D(Gi)-mCherry, 2.2 - 2.4 x10^13^ GC/mL) or mCherry control (n = 24, 13M/11F; listed above) into the BF. 400 nL of each virus was infused at target regions using a blunt-edged 33-gauge needle (Nanofil) at a rate of 100 nL/minute (World Precision Instruments), and left undisturbed for at least 10 minutes to allow for viral diffusion. Rats were given at least 3 weeks of viral incubation time before chemogenetic testing.

### Limited access drinking tests

One day following IAA and at least 2 weeks following surgery, all LTA and STA rats underwent daily 2-BC sessions with 15% ethanol to taper them to the final test length of 30 minutes, comprising three 12-16 hour overnight sessions and two 2-hour sessions, prior to receiving five consecutive ‘baseline’ 30-minute sessions (Fig. 1A). During the final three baseline sessions, animals were habituated to saline injections and a 30-minute post-injection wait period in test cages prior to ethanol access. Following baseline sessions, all LTA and STA rats received the chemogenetic ligand deschloroclozapine (DCZ, 0.1 mg/kg; Hello Bio) 30 minutes prior to a 30-minute 2-BC session with 15% ethanol and water. Following two re-baseline days with unadulterated 15% ethanol and saline injections, LTA and STA rats underwent additional DCZ testing sessions with increasing concentrations of quinine hydrochloride dihydrate (Sigma-Aldrich), consisting of either 45 (113 µM), 90 (226 uM), and 180 mg/L quinine (452 µM) or 15 (38 µM), 45, and 90 mg/L quinine. The results of 45 and 90 mg/L are reported here. These test sessions occurred every other day, and were interleaved with access to unadulterated ethanol following saline injections. Animals next underwent testing with 10% sucrose, 1-2 weeks following the final ethanol session. Animals received one overnight access to reduce neophobia, and three 30-minute baseline sucrose 2-BC sessions with saline injections prior to one DCZ-paired exposure to unadulterated 10% sucrose. Following two re-baseline sessions, rats received DCZ prior to access to sucrose with 45 mg/L quinine. To directly probe the effects of DCZ on drinking behavior in the absence of designer receptors a separate group of animals that did not undergo surgery were tested for ethanol consumption with counterbalanced injections of DCZ and saline. These rats underwent identical EtOH tapering and 30-minute baseline procedures to LTA and STA rats prior to the test sessions.

**Figure 1.**
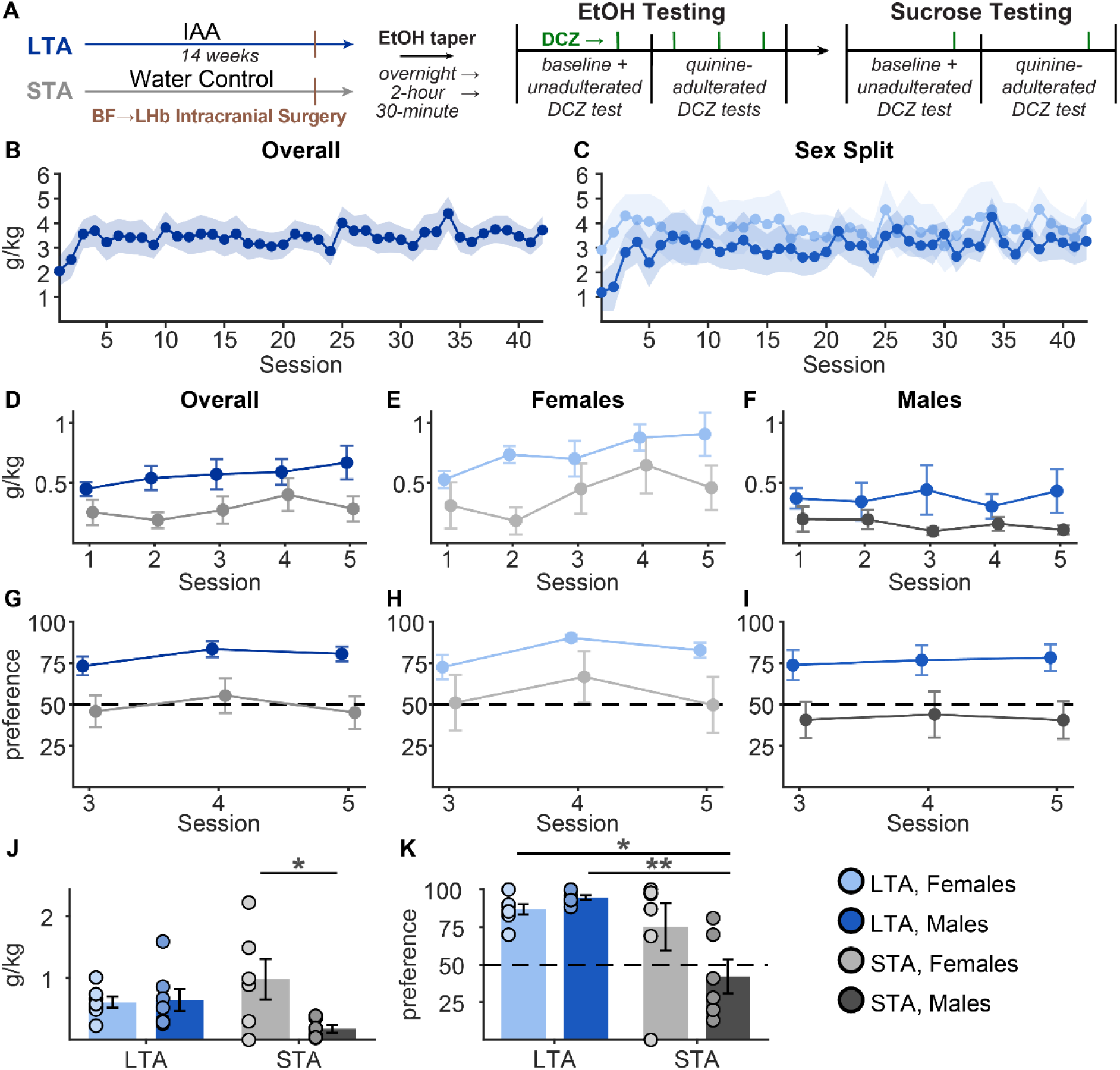
Long-term intermittent access to ethanol results in increased ethanol consumption and preference. (A) Experimental timeline. (B) Ethanol consumption (g/kg) during LTA in all animals (n=14), and (C) split by females (light blue, n=7) and males (medium blue, n=7). (D) Baseline 30-minute intake in LTA animals (blue) versus STA animals (grey; n = 12; 6F/6M) animals overall, and separately in (E) females and (F) males. (G-I) Ethanol preference measured during the last 3 days of baseline testing. (J) Ethanol consumption (g/kg) following DCZ. (K) Ethanol preference following DCZ. *p< 0.05, **p<0.01. Pairwise comparisons of estimated marginal means were conducted only following the identification of a significant interaction effect. Data are presented as mean ± SEM.

To assess blood ethanol concentrations (BECs) during from 30-minute drinking tests, 1-3 weeks after the end of behavioral testing 22 LTA (11F/11M) and 28 STA (14F/14M) rats were given access to 15% ethanol for 30 minutes and blood was collected via tail vein nick in heparinized capillary tubes. Blood samples were centrifuged at 10,000 RCF for 10 minutes at 4°C. Plasma was then aspirated and stored at -80°C until analysis. BECs were assessed using an AM1 Alcohol Analyser (Analox). BECs reliably reflected consumption following 30-minute ethanol intake (Fig. S1; R^2^ = 0.32, p < 0.001).

### Perfusion and tissue sectioning

Animals were deeply anesthetized with pentobarbital and underwent intracardial perfusion with 4% paraformaldehyde (PFA). Brains were extracted, post-fixed in PFA for 12-24 hours, and cryopreserved in 20% sucrose for at least 48 hours. Brains were then sectioned at 40 µM using either a cryostat (Leica; CM1860 UV) or microtome (Thermo Scientific; Microm HM450) and transferred to storage in 30% sucrose.

### Validation of virus placement in basal forebrain and Substance P immunohistochemistry

To assess the location and extent of viral expression, BF slices were stained for Substance P to illuminate the VP (Napier et al., 1991). 4 slices per animal were blocked in 10% normal donkey serum (NDS; Abcam), incubated for 12-16 hours (4°C) in rabbit anti-Substance P primary antibody (Immunostar, #20064; 1:2000), incubated for 2-3 hours (RT) in AlexaFluor 488 goat anti-rabbit secondary antibody (Invitrogen, #A-11008; 1:200), mounted, and coverslipped with DAPI-infused media (Vectashield). Images of the BF region were captured at 20x magnification with a 2x digital zoom using a Widefield Fluorescence Microscope (Keyence; BZ-X710) in DAPI, FITC (Substance P), and TRITC (mCherry) channels, with the same exposure settings in each channel across images. Using the 2 representative images exhibiting the greatest expression levels from each animal, cell bodies were counted using a custom semi-automated FIJI (RRID: SCR_002285) processing pipeline. mCherry-positive cells were assigned using the multi-point selection tool, and cell numbers for each hemisphere for each animal were averaged across images. Rats averaging ≤3 mCherry-positive cell bodies across representative 40 µm sections for either hemisphere were excluded from further analysis: 2/18 LTA hM3Dq rats (1/F/1M) and 14/34 STA Gi rats (4F/10M) were excluded. To assess subregional localization of viral expression, we employed a collection of open-source software packages based on the QUINT workflow (Yates et al., 2019). Atlas registration was performed on one representative 20x BF image from each hemisphere of each animal. Nutil (RRID: SCR_017183; Groeneboom et al., 2020) software was used to resize images, the QuickNII FileBuilder package (RRID: SCR_016854; Puchades et al., 2019) was used to transform images into XML files, and QuickNII base software was used to align slices to the CBWJ13 MR-histology rat atlas (Calabrese et al., 2013). Once aligned, VisuAlign (RRID: SCR_017978; Puchades et al., 2019) was used to manually warp atlas JSON files to fit over images. Finally, cell body multi-point selection ROIs from previous steps (see above) were re-aligned for each slice in FIJI, and cells within the BF were subregionally segmented based on overlaid atlas-derived boundaries.

### Assessment of viral expression in lateral habenula

Slices from LHb were stained for mCherry to amplify terminal expression using the procedures described above, using mouse anti-mCherry primary antibody (Living Colors, #632543; 1:1000), and AlexaFluor 555 goat anti-mouse secondary antibody (Invitrogen, #A21422; 1:200). All slices were imaged bilaterally at 10x with a Keyence in DAPI and TRITC (mCherry) channels. Axonal fluorescence was characterized with custom code in FIJI. LHb boundaries were hand drawn using the polygon selection tool, and fluorescence was quantified within the LHb and equivalently-sized loci within dorsal thalamic regions. LHb fluorescence in each hemisphere was normalized by subtracting dorsal thalamic signal to control for background fluorescence. Correlations between expression in the BF and LHb in each hemisphere were pooled across hM3Dq and hM4Di groups. One hM4Di male was excluded from correlation analysis due to outlier-level BF expression values (i.e., exceeding 3.5 SD above mean); this animal was retained for all behavioral analyses.

### Functional validation using cFos

To assess the functional impact of DCZ, rats expressing hM3Dq received either DCZ (0.1 mg/kg; 5F/4M) or saline vehicle (4F/4M) 60 minutes prior to perfusion. BF slices (2-4 per rat) were stained as described above using rabbit anti-cFos primary antibody (clone 9F6, Cell Signaling Technology, #2250S; 1:2500) and AlexaFluor 488 goat anti-rabbit secondary antibody (1:250). Images were captured in DAPI, FITC (cFos), and TRITC (mCherry) channels using an AXR confocal microscope (Nikon) with a 20x objective lens and 2x digital zoom. Colocalization of cFos and mCherry was measured for each cell with custom code in FIJI. Borders were hand drawn around BF cells expressing mCherry and superimposed with the FITC channel to generate a mixed color image. The Color Pixel Counter plugin (Ben Pichette) was used to calculate the number of pixels in the cFos channel (green), mCherry channel (red), and cFos/mCherry colocalization (green/red overlap, projected as yellow). Somatic colocalization score was calculated as the number of cFos/mCherry colocalized pixels divided by total mCherry pixels. Based on cumulative density distributions, we chose 20% overlap of cFos and mCherry as a cutoff for cFos-positive cells. We also assessed the impact of DCZ on VP and LHb cFos in control animals that did not undergo surgery. After injecting either DCZ or saline 60 minutes prior to sacrifice, we conducted IHC using rabbit anti-cFos primary antibody and AlexaFluor 555 donkey anti-rabbit secondary antibody (Invitrogen, #A31572; 1:250). All slices were imaged bilaterally at 20x with a Keyence in DAPI and TRITC channels. Equivalent borders were placed within the VP subregion of the BF and the LHb, and cFos positive cells were counted automatically within these borders using a custom FIJI script. Cell counts were averaged across each hemisphere for 2 images per animal (i.e., 4 total sections per animal) and normalized over ROI area.

### Statistical analysis

Data was processed and analyzed using RStudio and MATLAB. Linear mixed-effects models were used to examine the effects of experimental manipulations, with fixed effects for session, viral condition, and their interaction, with rat identify as a random effect. Age was also included as a random effect in analysis of STA animals. Sex was included as an additional fixed effect when relevant. When significant main effects or interactions were detected, pairwise contrasts of estimated marginal means were conducted with Sidak adjustment for multiple comparisons. Custom FIJI processing scripts are publicly available (doi.org/10.5281/zenodo.22820342) as well as tutorials describing our use of the QUINT workflow (doi.org/10.5281/zenodo.20630714).

## RESULTS

### Ethanol consumption during long-term intermittent access

In this study, we used a long-term intermittent access (LTA) model to promote ethanol drinking. Rats had access to 15% ethanol for 24-hour periods, 3 times per week for 14 weeks, consistent with IAA 2-BC procedures utilizing 20% ethanol (Simms et al., 2008; Wise, 1973). Prior to any exclusions, we observed that animals increased consumption across sessions (Fig. S2A; main effect of session: F(23,1150) = 2.44, p < 0.001) and that females consumed more than males (Fig. S2B; main effect of sex: F(1,50) = 4.32, p = 0.043; session by sex interaction, F(23,1150) = 0.78, p = 0.76). Starting at 8 weeks, animals were excluded from further study if they did not meet the following criteria: (1) overall average of >1.5 g/kg intake and (2) a rolling average of >1.5 g/kg intake across the 8 preceding sessions. Based on these criteria, 7/23 females and 11/27 males were excluded from subsequent testing, yielding 32 rats (16F/16M) for the testing phase. Following exclusions, animals that met criteria consumed similar amounts of ethanol across the entire 14 weeks (Fig. 1B; main effect of session: F(41,573) = 1.18, p = 0.21), with no significant effect of sex (Fig. 1C; main effect of sex: F(1,14) = 0.65, p = 0.43; session by sex interaction: F(41,573) = 0.65, p = 0.96). A separate group of rats (short-term access; STA) received a second water bottle during the IAA period, and, along with LTA rats, underwent five tapering sessions with access to 15% ethanol (three overnight sessions and two 2-hour sessions; Fig. 1A) prior to 30-minute baseline consumption and testing.

### Long-term intermittent access to 15% ethanol results in increased ethanol consumption and preference

We first analyzed whether LTA impacted ethanol intake in rats that received a control virus. Both LTA and STA rats underwent five baseline ethanol consumption sessions, prior to a test session with DCZ for later comparison to groups expressing designer receptors. During baseline sessions, LTA rats consumed more than STA rats (Fig. 1D; main effect of LTA: F(1,26) = 5.25, p = 0.030). When sex was included as a factor, we found that females consumed more than males (Fig. 1D-F; F(1,26) = 8.86, p = 0.006), but that sex did not interact with LTA (F(1,26) = 0.28, p = 0.60). We also found an interaction between sex and session (Fig. 1D-F; F(4,104) = 3.58, p = 0.009) that was driven by a significant effect of session in females (Fig. 1E; F(4,52) = 4.3, p = 0.004) but not in males (Fig. 1F; F(4,52) = 0.17, p = 0.95), suggesting females escalated intake over these 5 sessions to a greater extent than males. We also observed trends toward an effect of LTA in both females (Fig. 1E; main effect of LTA: F(1,13) = 4.48, p = 0.054) and males (Fig. 1F; F(1,13) = 2.7, p = 0.12). In addition to consuming more ethanol, LTA rats had a greater ethanol preference than STA animals (Fig. 1G; main effect of LTA: F(1,26) = 11.23, p = 0.003). When we analyzed preference separately for each sex, we observed a trend towards an increase in LTA females (Fig. 1H; main effect of LTA: F(1,13) = 3.52, p = 0.083) and a significant increase in LTA males (Fig. 1I; F(1,13) = 10.17, p = 0.007). Overall, LTA rats exhibited elevated ethanol intake and preference.

During the DCZ ethanol session, we observed a significant interaction between LTA and sex for consumption (Fig. 1J; F(1,22) = 4.96, p = 0.036) and preference (Fig. 1K; F(1,22) = 4.99, p = 0.036): pairwise comparisons indicated that STA females consumed more ethanol than STA males (Fig. 1J; p = 0.048), while LTA males and females had a higher ethanol preference than STA males (Fig. 1K; p = 0.003 and p = 0.013, respectively). Differences between LTA and STA females in ethanol intake during the DCZ day were not significant (Fig. 1J; p = 0.68), nor were differences between LTA females and STA males (p = 0.54). Elevated consumption and preference in STA females relative to STA males could reflect the baseline escalation in consumption we saw in females but not males or could reflect an off-target effect of DCZ. To address this question, we ran additional tests to determine whether DCZ produced non-selective effects on ethanol drinking or preference in a separate set of animals that did not undergo surgery. These animals received counterbalanced tests with DCZ or saline following identical tapering and baseline ethanol sessions to that STA animals received. In these animals we did not see a significant effect of DCZ on ethanol drinking (Fig. S3A; main effect of DCZ: F(1,14) = 1.8, p = 0.20) or preference (Fig. S3B; F(1,14) = 0.12, p = 0.73). When analyzing with sex as an additional factor, we found no main effect of sex or interaction between DCZ and sex for drinking (Fig. S3C-D; main effect of sex, F(1,14) = 0.57, p = 0.46; DCZ by sex interaction, F(1,14) = 0.01, p = 0.94) or preference (Fig. S3E-F; main effect of sex, F(1,14) = 0.02, p = 0.90; DCZ by sex interaction, F(1,14) = 0.12, p = 0.74). Therefore, the differences we found in LTA and STA animals likely do not reflect a non-specific effect of DCZ.

### Limited effect of long-term intermittent access on aversion-resistant ethanol intake-related behaviors

To examine aversion-resistant drinking, we compared consumption during the DCZ-paired 30-minute ethanol session to 30-minute test sessions with quinine-adulterated (45 and 90 mg/L) ethanol, which were also preceded by DCZ injections. Unexpectedly, we did not observe an impact of LTA (quinine concentration by LTA interaction, F(2,52) = 0.39, p = 0.68), only observing a significant main effect of quinine concentration (Fig. 2A; F(2,52) = 15.35, p < 0.001). When we included sex as a factor in our analysis, we observed a main effect of sex (F(1,26) = 7.5, p = 0.011), as well as an interaction between LTA and sex (F(1,26) = 4.5, p = 0.044), and quinine concentration, LTA, and sex (F(2,52) = 4.95, p = 0.011), indicating a sex-dependent difference in drinking between groups that varied based on quinine concentration. In females, we did not see a significant effect of LTA (Fig. 2B; F(1,13) = 1.64, p = 0.22) nor an interaction between LTA and quinine concentration (Fig. 2B; F(2,26) = 1.3, p = 0.29); however, in males, we observed a significant interaction between LTA and quinine concentration (Fig. 2C; F(2,26) = 7.42, p = 0.0028): both 45 and 90 mg/L quinine reduced ethanol consumption in LTA males (p = 0.010 and < 0.001, respectively), but not in STA males. This effect was largely driven by reduced consumption of unadulterated ethanol in STA males, which was significantly lower than LTA males (Fig. 2C; p = 0.037), resulting in a floor effect that prevented further reductions in consumption.

**Figure 2.**
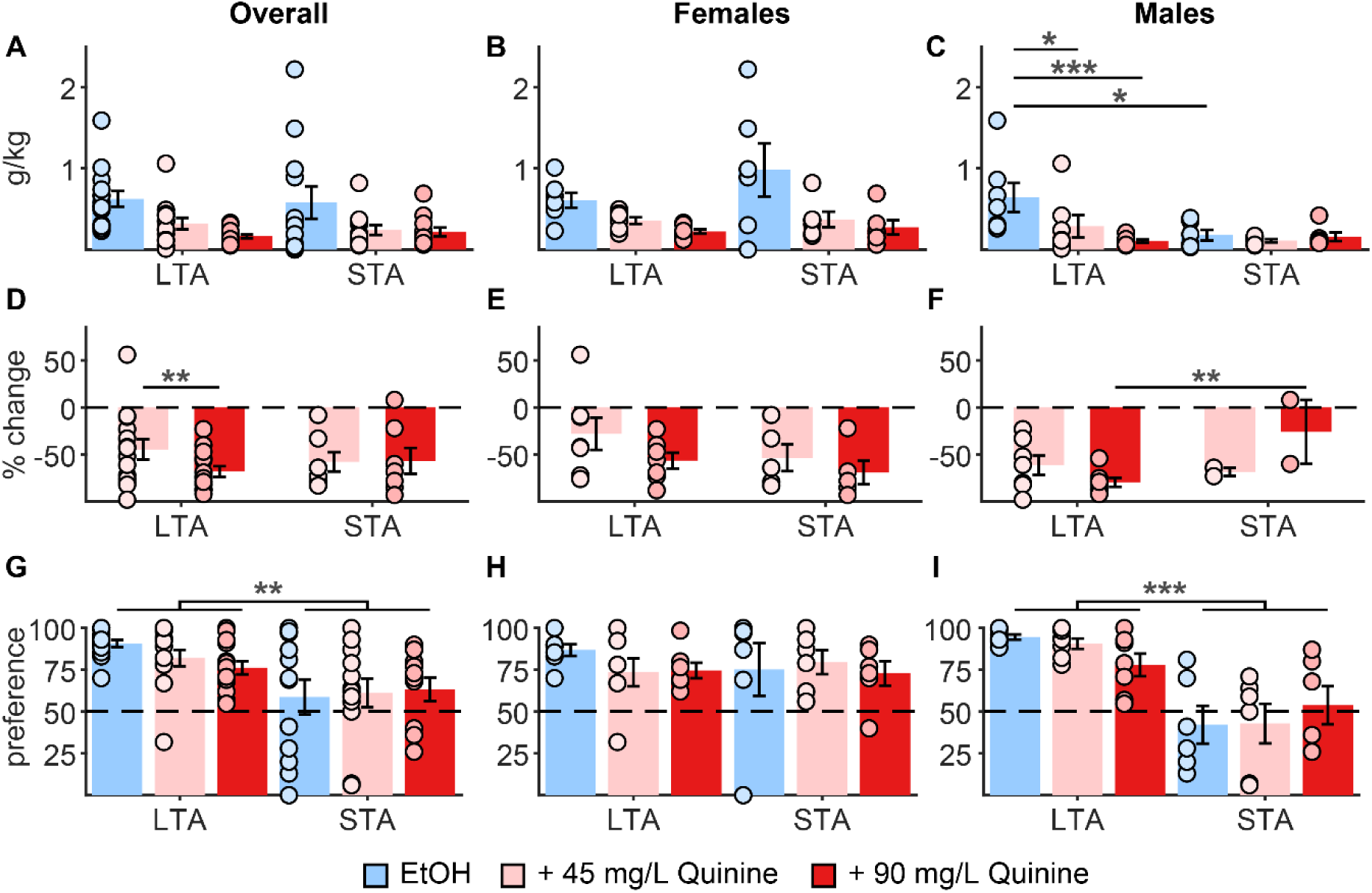
Long-term intermittent access promotes ethanol consumption and ethanol preference in the presence and absence of negative consequences in males. (A) DCZ-paired g/kg ethanol intake both with and without quinine for LTA (n=14; 7F/7M) and STA animals (n=12; 6F/6M) overall, and separately in (B) females and (C) males. (D) Percent change in ethanol intake following quinine adulteration overall, and separately in (E) females and (F) males (light blue, unadulterated 15% ethanol; pink, 45 mg/L adulterated 15% ethanol, medium red, 90 mg/L adulterated 15% ethanol). (G) DCZ-paired ethanol preference overall, and separately in (H) females and (I) males. *p< 0.05, **p<0.01, ***p<0.001. Pairwise comparisons of estimated marginal means were conducted only following the identification of a significant interaction effect. Data are presented as mean ± SEM.

To better account for baseline differences in consumption and avoid floor effects, we excluded animals with baseline ethanol intake below 0.2 g/kg (1F/4M STA animals) and calculated the % change from consumption of unadulterated ethanol (Fig. 2D-F). We observed a significant main effect of quinine concentration (F(1,42) = 4.78, p = 0.034) and a significant interaction between LTA and quinine concentration (F(1,42) = 5.54, p = 0.023) but no main effect of LTA (F(1,21.94) = 0.01, p = 0.93): LTA animals diminished consumption to greater levels as quinine concentrations progressed (p = 0.002), while STA animals suppressed similarly at both concentrations (Fig. 2D). When adding sex as a factor, we saw additional significant interactions between quinine concentration and sex (F(1,42) = 12.97, p < 0.001) as well as quinine concentration, LTA, and sex (F(1,42) = 6.29, p = 0.016). While we only observed a significant main effect of quinine concentration in females (Fig. 2E; F(1,24) = 24.58, p < 0.001), we observed an interaction between LTA and quinine concentration in males (Fig. 2F; F(1,18) = 10.39, p = 0.005), observing greater suppression of consumption in LTA males at the 90 mg/L condition. However, this effect should be interpreted cautiously, as it reflects data from only two STA animals with relatively high baseline ethanol intake, excluding the majority of STA males that consumed minimal ethanol.

For quinine-adulterated ethanol preference, we saw a significant main effect of LTA (Fig. 2G; F(1,26) = 11.3, p = 0.002) without an interaction between LTA and quinine concentration (F(2,52) = 1.78, p = 0.18), indicating LTA animals exhibited greater preference for ethanol across these sessions. When analyzing with sex as a factor, we saw a strong trend towards a main effect of sex (Fig. 2G; F(1,26) = 4.03, p = 0.055) and an interaction between LTA and sex (F(1,26) = 15.17, p = <0.001). Upon analyzing males and females separately, we found no significant effect of LTA (F(1, 39) = 0.15, p = 0.70) on ethanol preference in females, but LTA males had greater ethanol preference than STA males (Fig. 2I; F(1,13) = 26.77, p = <0.001).

Notably, we did not observe a significant effect of quinine in any of our analyses examining ethanol preference. Overall, while LTA did not promote a strong aversion-resistant drinking phenotype, it did enhance ethanol preference both with and without quinine in males.

### Quinine adulteration reduces sucrose consumption similarly across groups

We next assessed the impact of quinine adulteration (45 mg/L) on sucrose consumption. Male and female LTA and STA animals consumed similar g/kg of 10% sucrose across baseline sessions (Fig. 3A; main effect of session: F(2,51.59) = 0.22, p = 0.81; main effect of LTA: F(1,26.24) = 0.57, p = 0.46; main effect of sex: F(1,26.26) = 1.84, p = 0.19). Animals then received DCZ before test sessions with 10% sucrose adulterated with either 0 or 45 mg/L quinine. As expected, quinine reduced g/kg sucrose consumption (Fig. 3C; F(1,26) = 80.96, p < 0.001) with no effect of LTA (F(1,26) = 0.14, p = 0.71). Because we saw a significant main effect of sex (F(1,26) = 5.47, p = 0.027), and an LTA by sex interaction (F(1,26) = 5.52, p = 0.027), we analyzed each sex independently. We found no significant effect of LTA in females (Fig. 3D; F(1,13) = 1.3, p = 0.28), but in males we found a main effect of LTA (Fig. 3E; F(1,13) = 6.5, p = 0.024) and a quinine by LTA interaction (Fig. 3E; F(1,13) = 9.32, p = 0.009): LTA and STA males both drank less quinine-adulterated sucrose, but STA males drank significantly less unadulterated sucrose than LTA males (p = 0.006). When we assessed % change from baseline, excluding animals consuming <3g of sucrose to avoid a floor effect (2 STA animals, 1M/1F) we did not see a significant difference between LTA and STA animals (Fig. 3F; t(22) = - 0.60, p = 0.56).

**Figure 3.**
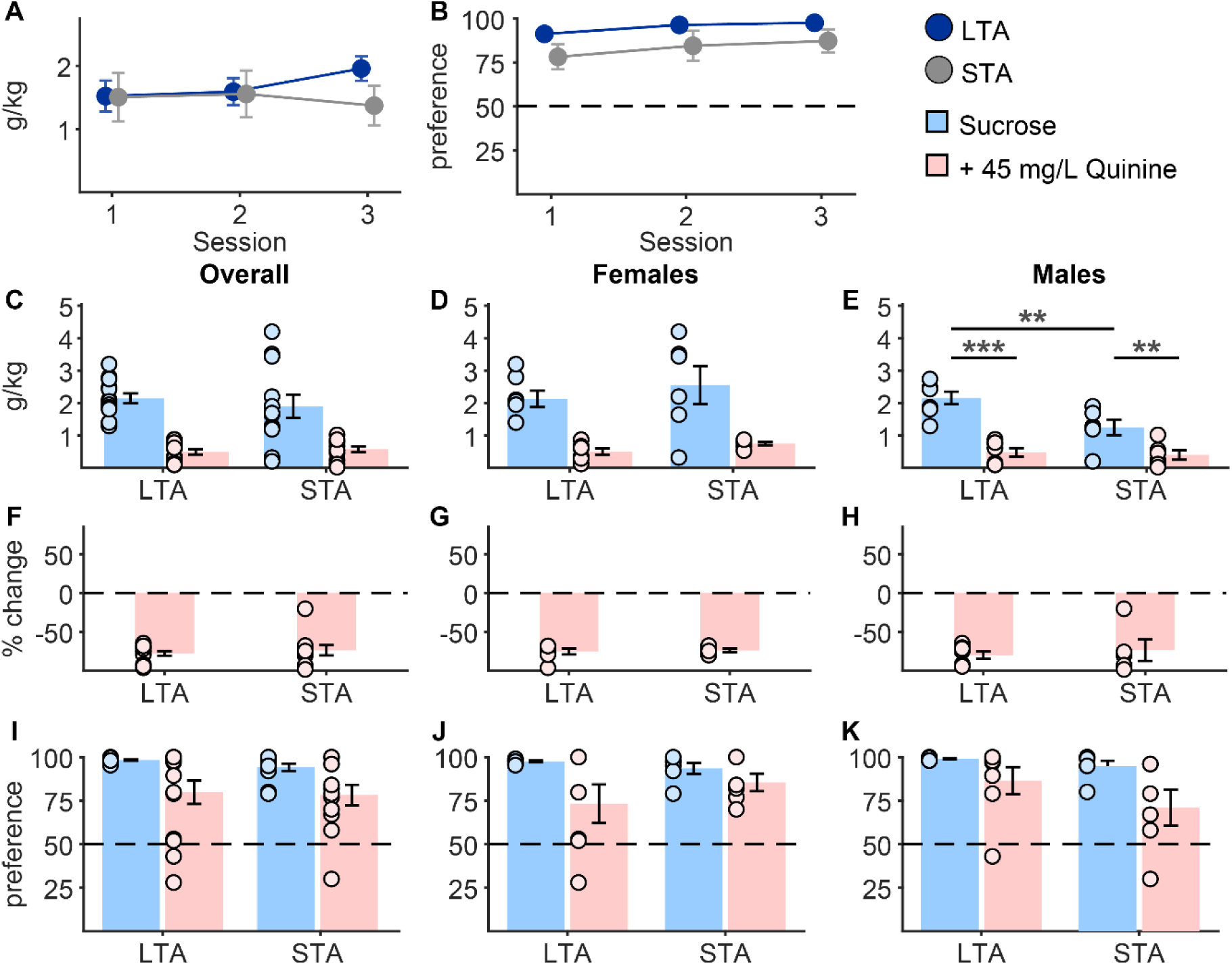
Quinine adulteration reduces sucrose consumption similarly across groups. (A-B) 3-day baseline 30-minute sucrose intake and preference. (A) Baseline g/kg sucrose intake and (B) preference overall between LTA (blue; n=14; 7F/7M) and STA (grey; n=12; 6F/6M) rats. (C) DCZ-paired g/kg sucrose intake both with and without quinine overall, and separately in (D) females and (E) males (light blue, unadulterated 10% sucrose; pink, 45 mg/L adulterated 10% sucrose). (F) Percent change in sucrose intake following quinine adulteration overall, and separately in (G) females and (H) males. (I) DCZ-paired sucrose preference overall, and separately in (J) females and (K) males. **p<0.01, ***p<0.001. Pairwise comparisons of estimated marginal means were conducted only following the identification of a significant interaction effect. Data are presented as mean ± SEM.

When we assessed baseline sucrose preference, we found trends towards a main effect of LTA (Fig. 3B; F(1,26.67) = 3.55, p = 0.070), suggesting that LTA may have marginally enhanced baseline sucrose preference. During DCZ-paired sessions, we found a significant effect of quinine (F(1,52) = 14.76, p < 0.001) but did not see a significant effect of LTA (Fig. 3I; F(1,52) = 0.41, p = 0.53) or quinine by LTA interaction (F(1, 52) = 0.08, p = 0.78). We did not see a main effect of sex (F(1, 52) = 0.01, p = 0.93), but we saw a trend towards an LTA by sex interaction (F(1, 52) = 2.67, p = 0.11) and thus chose to analyze preference data separated by sex. While we found an impact of quinine for both sexes (females, F(1, 13) = 7.37, p = 0.018; males, F(1, 26) = 9.24, p = 0.005) we saw no impact of LTA (Fig. 3J-K; main effect of LTA: females, F(1,13) = 0.44, p = 0.52; males Fig. 3K; F(1,26) = 2.69, p = 0.11) or quinine by LTA interactions (females, F(1, 13) = 1.87, p = 0.19; males, F(1, 26) = 0.89, p = 0.35), indicating the quinine similarly reduced sucrose preference across each group.

### Expression and functional validation of designer receptors in basal forebrain neurons projecting to the lateral habenula

Next, we assessed the impact of chemogenetic activation or inhibition of BF-to-LHb projections on ethanol and natural reward consummatory behaviors. After LTA or STA exposure, rats received virus for cre-dependent expression of hM3Dq (Gq), hM4Di (Gi), or control fluorophore (mCherry) in the BF, and a retrograde virus for cre-recombinase in the LHb. Animals expressing fewer than 3 mCherry-labeled neurons in either hemisphere averaged across BF sections were excluded from further analysis. We then analyzed localization of viral expression via manual atlas alignment processing based on the QUINT protocol (Yates et al., 2019), aligning one image per hemisphere per rat in each experimental group to the MRH rat atlas (Fig. S4; Calabrese et al., 2013). In our atlas-aligned sections, we observed approximately 17.1 ± 0.2 mCherry-labeled BF-to-LHb neurons per hemisphere across all experimental animals (Figure 4A-B). The majority of viral expression was observed to be localized within the VP across all groups (Figure 4A-B; 53.5 ± 3.9%). We also found that LHb mCherry fluorescence intensity correlated strongly with mCherry-labeled cell count within the BF of each hemisphere (Fig. 4C-D; R^2^ = 0.23, p < 0.001). To assess the functional impact of DCZ, we assessed DCZ-evoked cFos in the BF of rats expressing Gq virus (Fig. 4E). We found that DCZ enhanced cFos colocalization with virus (73.7 ± 8.4%) relative to saline (16.2 ± 4.8%) (Fig. 4F; t(15) = 5.77, p < .001). We found no effect of sex (F(1, 13) = 1.21, p = 0.29) or interaction between sex and DCZ (F(1, 13) = 2.08, p = 0.17). In surgery-free control animals that were tested for the behavioral impact of DCZ, we also assessed the impact of DCZ on cFos and found that in the absence of expression of designer receptors DCZ did not alter cFos expression in the VP (Fig. S5; main effect of DCZ, t(12) = 0.83, p = 0.42; DCZ by sex interaction, F(1,10) = 0.02, p = 0.88), or in the LHb (data not shown; main effect of DCZ, t(12) = 1.28, p = 0.22; DCZ by sex interaction, F(1,10) = 1.15, p = 0.31).

**Figure 4.**
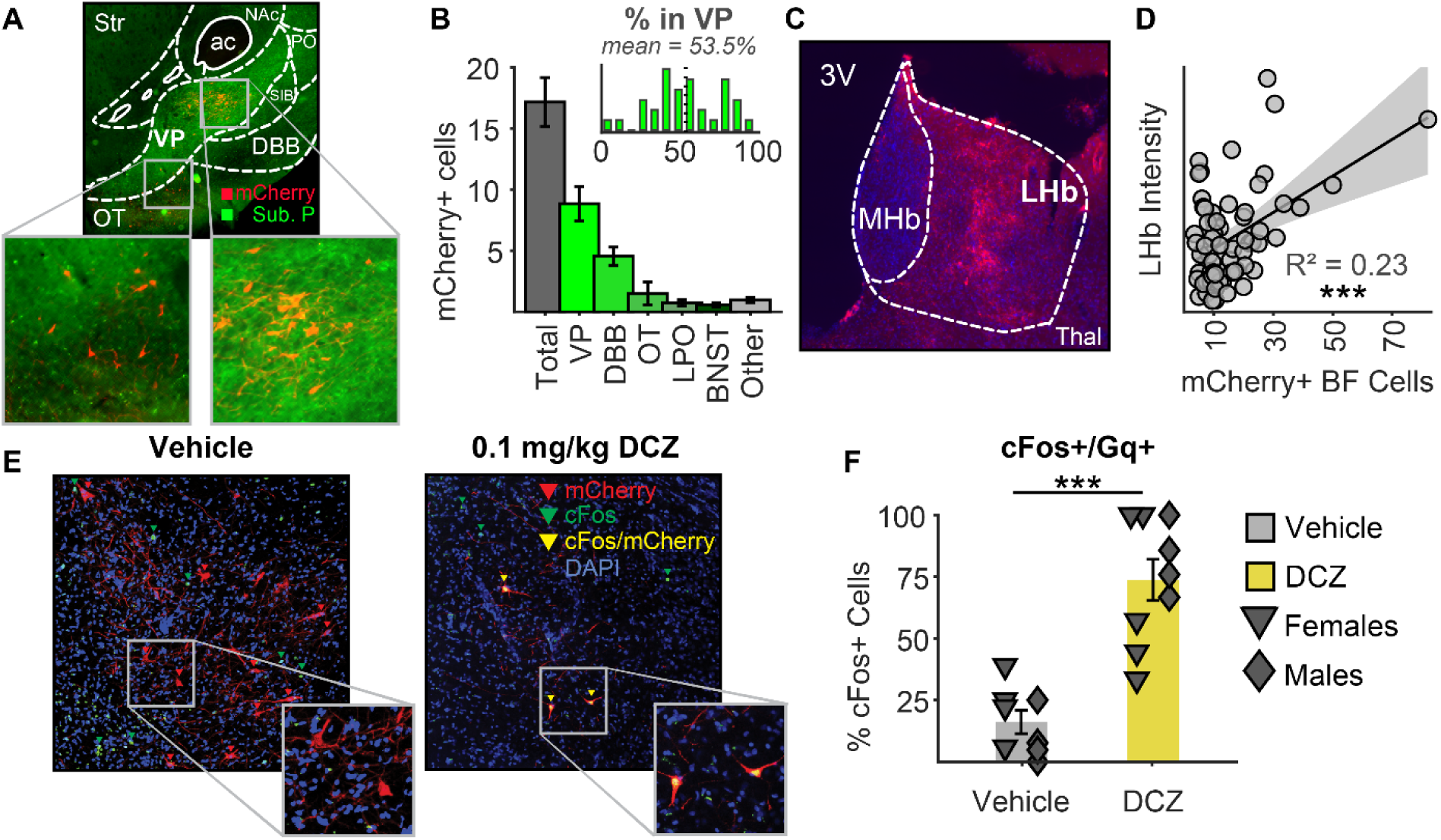
Expression and functional validation of designer receptors in BF neurons projecting to the LHb. (A) diagram of example mCherry-positive cell body expression in the basal forebrain following substance P immunostain. (B) breakdown of subregional expression patterns in the BF and histogram of percent expression within ventral pallidum. VP = ventral pallidum; DBB = diagonal band of Broca; OT = olfactory tubercles; LPO = lateral preoptic area; BNST = basal nucleus of the stria terminalis; SIB = substantia innominata; Str = striatum; NAc = nucleus accumbens; ac = anterior commissure. (C) diagram of example mCherry-positive axonal expression in the lateral habenula following mCherry amplification immunostain. LHb = lateral habenula; MHb = medial habenula; Thal = thalamus; 3V = 3^rd^ ventricle. (D) normalized fluorescence intensity in the LHb correlated to # mCherry-expressing neurons in each hemisphere. (E) diagram of example cFos colocalization in Gq-expressing neurons following either DCZ or saline pre-injection 60 minutes prior to sacrifice. (F) quantification of cFos/Gq colocalization following DCZ (yellow) or vehicle (grey) pre-injection, split by females (triangles) and males (diamonds). ***p<0.001. Data are presented as mean ± SEM.

### Chemogenetic activation of BF-to-LHb neurons has opposing effects on ethanol consumption patterns in males and females

We originally hypothesized that BF-to-LHb activation would diminish aversion-resistant ethanol consumption in LTA rats. However, we did not find strong evidence for aversion-resistant drinking in LTA rats expressing control virus (Fig. 2). Nevertheless, we analyzed these data to see if activation of this pathway altered ethanol drinking behavior in the presence or absence of quinine adulteration. Across IAA, LTA animals drank similarly both between viral groups (Fig. S2C; main effect of viral group: F(1,33) = 0.49, p = 0.49; session by viral group interaction: F(41,1352) = 0.66, p = 0.95) and between sexes (Fig. S2D-E; main effect of sex: F(1,33) = 1.74, p = 0.20; session by sex interaction: F(41,1352) = 0.76, p = 0.86). Importantly, we observed no significant effects of viral group on ethanol consumption (Fig. 5A; F(1,30) = 0.05, p = 0.83) or preference (Fig. 5D; F(1,30.98) = 0.01, p = 0.91) during baseline sessions. We found that LTA females consumed greater g/kg ethanol across baseline sessions than males (Fig. 5B-C; F(1,30) = 12.67, p = 0.001), similarly between viral group (sex by viral group interaction, F(1, 30) = 0.004, p = 0.95). Ethanol preference increased across baseline sessions (Fig. 5D; F(2,57.76) = 3.16, p = 0.050), with no impact of viral group (F(1,30.98) = 0.01, p = 0.91) or sex (F(1,31.13) = 2.14, p = 0.15).

**Figure 5.**
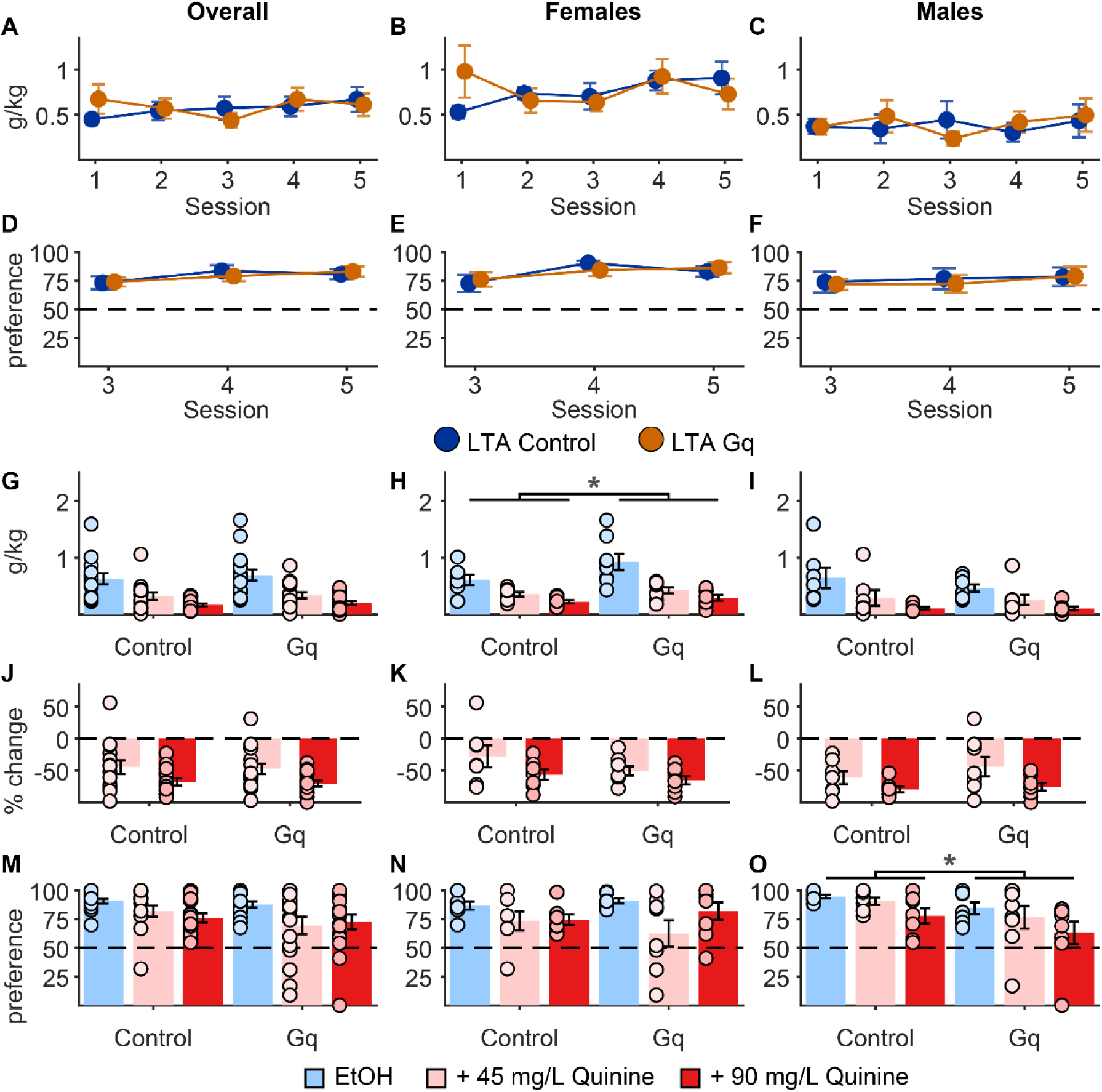
Chemogenetic activation of BF-to-LHb neurons has opposing effects on ethanol consumption patterns in males and females. (A) Baseline 30-minute intake in LTA control (dark blue; n=14; 7F/7M) and LTA Gq animals (orange; n = 16; 8F/8M) overall, and separately in (B) females and (C) males. (D-F) Ethanol preference measured during the last 3 days of baseline testing. (G) DCZ-paired g/kg ethanol intake both with and without quinine for LTA control (n=14; 7F/7M) and LTA Gq animals (n = 16; 8F/8M) overall, and separately in (H) females and (I) males (light blue, unadulterated 15% ethanol; pink, 45 mg/L adulterated 15% ethanol, medium red, 90 mg/L adulterated 15% ethanol). (J) Percent change in ethanol intake following quinine adulteration overall, and separately in (K) females and (L) males. (M) DCZ-paired ethanol preference overall, and separately in (N) females and (O) males. *p< 0.05. Data are presented as mean ± SEM.

When we analyzed DCZ sessions, we found that while quinine reduced ethanol consumption (Fig. 5G; main effect of quinine concentration, F(2,60) = 45.5, p < 0.001), there was no effect of Gq virus (F(1,30) = 0.29, p = 0.59), or interaction between viral group and quinine concentration (F(2,60) = 0.13, p = 0.88). Because females consumed more g/kg ethanol during DCZ sessions (main effect of sex: F(1, 30) = 6.33, p = 0.017) with a trend towards an interaction with viral group (viral group by sex interaction: F(1,30) = 3.01, p = 0.093, we analyzed these data separately for each sex. In females, we found that Gq animals consumed more ethanol (Fig. 5H; F(1,15) = 4.90, p = 0.043), regardless of quinine concentration (virus by quinine concentration interaction: F(2,30) = 2.0, p = 0.15). In males, we did not observe a significant effect of viral group (Fig. 5I; F(1,15) = 0.43, p = 0.52). When we analyzed % change, we found an effect of quinine concentration (Fig. 5J; F(1,30) = 21.48, p < 0.001), but no effect of viral group (Fig. 5J; main effect of viral group: F(1,30) = 0.09, p = 0.77; quinine concentration by viral group interaction: F(1,30) = 0, p = 0.99). While there were trends toward a main effect of sex (F(1,30) = 3.41, p = 0.075), and a viral group by sex interaction (F(1,30) = 2.52, p = 0.12) we did not see any viral group differences in the impact of quinine when separating by sex (Fig. 5K-L; main effect of viral group: females, F(1,15) = 1.65, p = 0.22; males, F(1,15) = 0.9, p = 0.36). These data indicate that while females increased overall g/kg consumption upon Gq activation, regardless of quinine adulteration, they suppressed intake to a similar extent as control females when considering individual baseline consumption.

In males and females combined, we also found that quinine reduced ethanol preference (Fig. 5M; F(2, 58.34) = 5.43, p = 0.007) regardless of viral group (main effect of viral group, F(1, 29.23) = 2.1, p = 0.16; quinine concentration by viral group interaction, F(1, 58.34) = 0.59, p = 0.56). Because the impact of quinine adulteration on ethanol preference varied by sex (Fig. 5M-O; quinine concentration by sex interaction: F(2, 57.78) = 3.36, p = 0.042) we next analyzed males and females separately. While quinine significantly reduced preference in females (Fig. 5N; F(2, 27.99) = 4.75, p = 0.017) we found no effect to Gq virus (F(1, 13.47) = 0, p = 0.99). In contrast, we found both a main effect of quinine concentration in males (Fig. 5O; F(2, 29.64) = 4.79, p = 0.016) and a significant effect of viral group (F(1, 15.13) = 4.62, p = 0.048) but no quinine concentration by viral group interaction (F(2, 29.64) = 0.09, p = 0.91), suggesting that males in the Gq group had lower preference for ethanol with and without quinine. These findings suggest that the effects of BF-to-LHb activation may vary by sex: activation potentiated ethanol consumption in females and reduced ethanol preference in males, all in a manner that did not depend on quinine concentration.

### BF-to-LHb activation may decrease sensitivity to quinine adulteration of sucrose

We next assessed whether BF-to-LHb activation impacted sucrose consumption. During baseline sessions, we observed no effects of viral group (Fig. 6A; F(1,30.16) = 0, p = 0.97) or sex (F(1, 30.16) = 0.52, p = 0.48) on sucrose intake. No effect of viral group was observed on baseline sucrose preference (Fig. 6B; F(1,32.35) = 0.84, p = 0.37). During DCZ sessions (Fig. 6C-E), we saw a significant main effect of quinine (F(1,24.59) = 94.99, p < 0.001), and a significant interaction between quinine and viral group (F(1,24.59) = 7.53, p = 0.011). This interaction was not explained by any *post hoc* comparisons (Fig. 6C). No significant sex differences were observed (Fig. 6D-E; F(1,25.18) = 2.51, p = 0.13). After excluding animals with low sucrose consumption (2 Gq animals, 1M/1F) we saw that BF-to-LHb activation significantly reduced the magnitude of quinine-evoked suppression (% change; t(25) = 3.16, p = 0.004). This effect was significant in females (t(11) = 2.73, p = 0.019) but not in males (t(12) = 2.00, p = 0.069). For sucrose preference, we found a main effect of quinine (Fig. 6I; F(1,29.97) = 9, p = 0.005), with a strong trend towards an interaction between quinine and viral group (F(1,29.97) = 3.83, p = 0.0598). Although this interaction did not reach statistical significance, given our *a priori* interest in the impact of BF-to-LHb activation on quinine sensitivity, we chose to examine pairwise comparisons to further characterize the pattern of effects. *Post hoc* comparisons showed a significant reduction in sucrose preference following quinine exposure amongst controls (Fig. 6I; p = 0.0095), but no significant difference amongst Gq animals following quinine adulteration (p = 0.92). These findings suggest BF-to-LHb activation reduced sensitivity to quinine adulteration.

**Figure 6.**
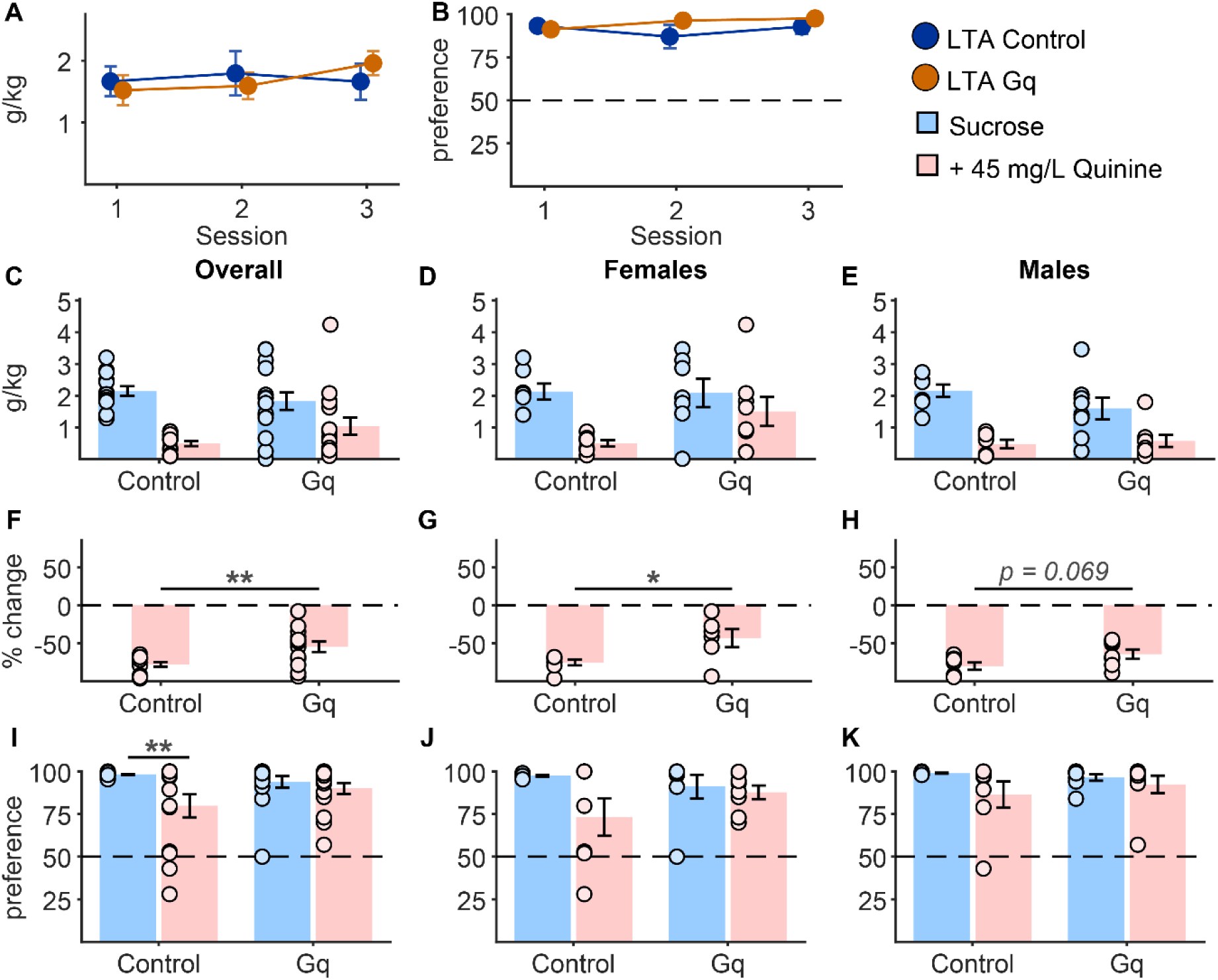
BF-to-LHb activation may decrease sensitivity to quinine adulteration of sucrose. (A) Baseline g/kg sucrose intake and (B) preference overall between LTA control (dark blue; n=14; 7F/7M) and LTA Gq (orange; n = 16; 8F/8M) rats. (C) DCZ-paired g/kg sucrose intake both with and without quinine overall, and separately in (D) females and (E) males (light blue, unadulterated 10% sucrose; pink, 45 mg/L adulterated 10% sucrose). (F) Percent change in sucrose intake following quinine adulteration overall, and separately in (G) females and (H) males. (I) DCZ-paired sucrose preference overall, and separately in (J) females and (K) males. *p< 0.05, **p<0.01. Data are presented as mean ± SEM.

### Chemogenetic inhibition of BF-to-LHb neurons does not impact ethanol consumption in animals with limited ethanol history

We next investigated whether chemogenetic inhibition of BF-to-LHb neurons impacts ethanol consumption in the presence and absence of quinine adulteration in STA rats. During baseline sessions, we found an interaction between session and viral group (Fig. 7A-C; F(4,175.11) = 2.96, p = 0.021) and session, viral group and sex (F(4,175.17) = 5.52, p < 0.001). Female Gi animals consumed more during the first session (p = 0.008) but not during subsequent baseline sessions (Fig. 7B; session by viral group interaction: F(4,88) = 5.16, p < 0.001). Baseline ethanol preference also varied based on session, viral group, and sex (Fig. 7D; 3-way interaction: F(2,85.58) = 5.44, p = 0.006). Preference in males was influenced by an interaction of session and virus (Fig. 7F; F(2,41.52) = 3.69, p = 0.033), but no pairwise differences explained these interactions. During DCZ test sessions, we saw a significant effect of quinine concentration (Fig. 7G; F(2,88) = 15.34, p < 0.001), but no effect of viral group (main effect: F(1,44) = 0.01, p = 0.92; interaction with quinine concentration, F(2, 88) = 0.46, p = 0.63). Females consumed more ethanol (Fig. 7H-I; F(1,44) = 4.18, p = 0.047), but we observed no effects of virus or interaction with quinine concentration in either females or males. Similarly, when we examined % change from baseline after exclusions (<0.2 g/kg baseline consumption: 8 control, 2F/6M; 7 Gi, 3F/4M), we found an effect of quinine concentration (Fig. 7J; F(2,58) = 27.49, p < 0.001), but no effect of viral group (main effect: F(1,29) = 0.21, p = 0.65; interaction with quinine concentration: F(2,58) = 0.1, p = 0.91). While we saw a trend towards an effect of sex (Fig. 7K-L; F(1,29) = 3.03, p = 0.092), we observed no interactions and no effects of viral group when analyzed separately. For ethanol preference, we observed no effects of quinine concentration (Fig. 7M; F(2,87.12) = 1, p = 0.37) or viral group (main effect of viral group: F(1,43.88) = 0.01, p = 0.94; quinine concentration by viral group interaction: F(2,87.12) = 1.31, p = 0.27). Females tended to have higher ethanol preference (Fig. 7N-O; F(1,44.01) = 3.7, p = 0.061), but no significant effects of viral group were found when sexes were analyzed separately (main effect of viral group, females: F(1,22) = 0.34, p = 0.57; males, F(1,22.01) = 0.04, p = 0.85; quinine concentration by viral group interaction, females: F(2,44) = 0.21, p = 0.81; males, F(2,43.26) = 1.51, p = 0.23). Overall BF-to-LHb inhibition did not influence ethanol intake or preference in the presence or absence of quinine adulteration.

**Figure 7.**
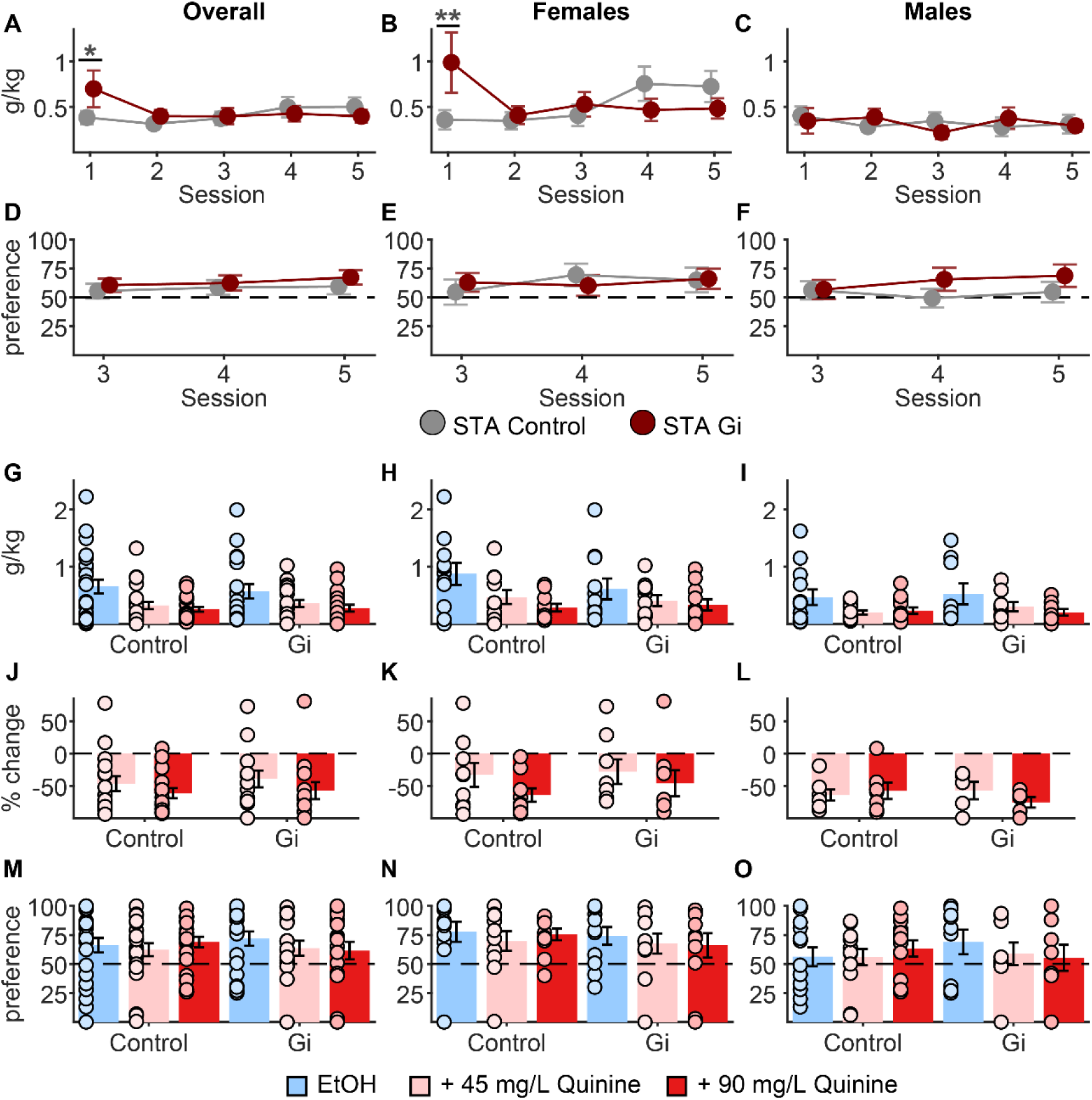
Chemogenetic inhibition of BF-to-LHb neurons does not impact ethanol consumption in animals with limited ethanol history. (A) Baseline 30-minute intake in STA control (grey; n = 24; 11F/13M) and STA Gi animals (dark red; n = 20; 11F/9M) overall, and separately in (B) females and (C) males. (D-F) Ethanol preference measured during the last 3 days of baseline testing. (G) DCZ-paired g/kg ethanol intake both with and without quinine for LTA control (n=14; 7F/7M) and LTA Gq animals (n = 16; 8F/8M) overall, and separately in (H) females and (I) males (light blue, unadulterated 15% ethanol; pink, 45 mg/L adulterated 15% ethanol, medium red, 90 mg/L adulterated 15% ethanol). (J) Percent change in ethanol intake following quinine adulteration overall, and separately in (K) females and (L) males. (M) DCZ-paired ethanol preference overall, and separately in (N) females and (O) males. *p< 0.05**, p<0.01. Pairwise comparisons of estimated marginal means were conducted only following the identification of a significant interaction effect. Data are presented as mean ± SEM.

### Chemogenetic inhibition of BF-to-LHb neurons does not impact sucrose consumption

Finally, we tested whether chemogenetic inhibition of BF-to-LHb neurons impacts sucrose intake. We did not observe any differences in baseline sucrose consumption based on viral group (Fig. 8A; F(1,43.28) = 0.77, p = 0.38) or sex (F(1,43.34) = 0.12, p = 0.73). While sucrose preference increased across baseline sessions (Fig. 8B; F(2,77.06) = 3.5, p = 0.035) this did not differ based on viral group (F(1,42.56) = 0.11, p = 0.74) or sex (F(1,42.23) = 0.41, p = 0.53). During DCZ sessions, quinine reduced sucrose consumption (Fig. 8C; F(1,44) = 68.22, p < 0.001) similarly between viral groups (quinine by viral group interaction: F(1,44) = 0, p = 0.99). We observed no effect of sex (Fig. 8D-E; F(1,42.72) = 1.68, p = 0.20). After exclusions (<3g baseline consumption: 2 control, 1F/1M; 1 Gi, 1F) we also found no effect of viral group on % change (Fig. 8F; t(38) = 0.58, p = 0.56). Quinine also reduced sucrose preference (Fig. 8I; F(1,44) = 19.67, p < 0.001) similarly between viral groups (quinine by viral group interaction: F(1,44) = 1.6, p = 0.21). While we observed a trend towards an interaction between sex and quinine (Fig. 8I-K; F(1,44) = 2.99, p = 0.091), no differences were observed between viral groups within each sex (quinine by viral group interaction, females: F(1,22) = 0.02, p = 0.9; males: F(1,22) = 1.84, p = 0.19). Thus, we found that inhibition of BF-to-LHb pathway did not impact sucrose consumption or preference.

**Figure 8.**
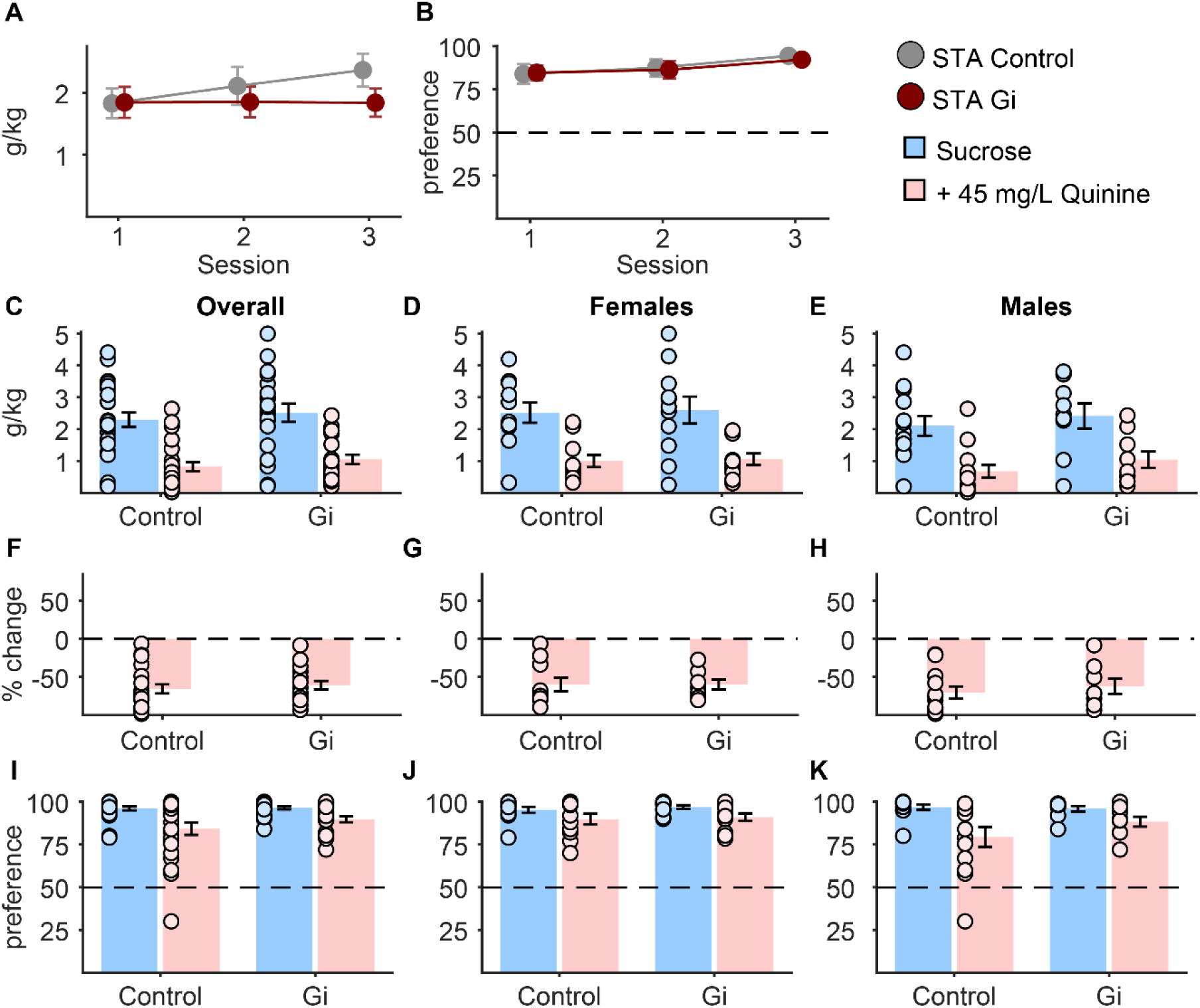
Chemogenetic inhibition of BF-to-LHb neurons does not impact basal or quinine-adulterated sucrose consumption. (A) Baseline g/kg sucrose intake and (B) preference overall between STA control (grey; n = 24; 11F/13M) and STA Gi (dark red; n = 20; 11F/9M) rats. (C) DCZ-paired g/kg sucrose intake both with and without quinine overall, and separately in (D) females and (E) males (light blue, unadulterated 10% sucrose; pink, 45 mg/L adulterated 10% sucrose). (F) Percent change in sucrose intake following quinine adulteration overall, and separately in (G) females and (H) males. (I) DCZ-paired sucrose preference overall, and separately in (J) females and (K) males. Data are presented as mean ± SEM.

## DISCUSSION

Here, we studied the effects of long-term intermittent access to alcohol and chemogenetic manipulation of BF-to-LHb projecting neurons on ethanol and natural reward intake. After long-term ethanol access, animals elevated ethanol drinking and preference. Upon quinine adulteration, we observed no evidence of aversion-resistant intake, although males showed a stronger preference for quinine-adulterated ethanol following long-term access. We found that chemogenetic activation of BF-to-LHb neurons has opposing effects on ethanol consumption patterns in males and females. Activation suppressed preference for ethanol with and without quinine in males. In contrast, chemogenetic activation of BF-to-LHb neurons increased consumption of ethanol in females. Activation also diminished suppression of quinine-adulterated sucrose similarly between sexes. Chemogenetic inhibition of BF-to-LHb neurons had no effect on ethanol or sucrose consumption, in the presence or absence of quinine.

### Absence of aversion-resistant ethanol drinking phenotype

Prior research has found that rats exposed to long-term IAA (Simms et al., 2008; Wise, 1973) demonstrate sustained motivation for ethanol, continuing seeking and intake despite the presence of quinine (Hopf et al., 2010; Seif et al., 2013, 2015). Despite receiving 14 weeks of intermittent ethanol access, we did not observe robust aversion-resistant ethanol intake in LTA rats relative to STA rats. This may be due to several differences between our protocol and that of prior work looking at aversion-resistant drinking following intermittent access. Of note, prior studies that documented aversion-resistant drinking after intermittent access used 20% ethanol in male Wistar rats, while we used 15% ethanol in Long-Evans rats of both sexes. IAA with 15% ethanol has been used to promote ethanol drinking in advance of cue conditioning in several studies (Carpio et al., 2022; Cofresí et al., 2017, 2018, 2019; LeCocq et al., 2022; Millan et al., 2015, 2017; Segal et al., 2022; Sood et al., 2026; Sparks et al., 2014) and promotes ethanol seeking behavior in male Long-Evans rats (Angelyn et al., 2021). Yet, the use of 15% ethanol IAA to promote aversion-resistant drinking has not been well characterized, which may have contributed to the lack of aversion-resistant drinking in our study. Nevertheless, g/kg ethanol intake during 15% ethanol IAA remained similar in Long-Evans rats in comparison to standard values observed in the field (Fig. 1 and S2; see Carnicella et al., 2014), suggesting that the use of 15% ethanol alone may not fully explain the absence of aversion-resistant drinking. To this end, other studies utilizing a similar LTA model in rats began quinine titrations at lower concentrations than used in our study. While Hopf et al. (2010) and Seif et al. (2013) established aversion-resistant drinking with 10 and 30 mg/L during various lengths of 2-BC ethanol, Hopf et al. (2010) also observed an even stronger aversion-resistance phenotype when testing with concentrations as high as 100 mg/L quinine, though they observed this in a progressive ratio task following LTA. Based on this evidence that aversion resistance can extend to higher quinine concentrations, both in these alternative behaviors as well as in preliminary data from our lab, the lowest concentration we tested in the present study was 45 mg/L. We may have observed aversion resistance if we had tested at lower quinine concentrations. Additionally, prior work incorporated repeated testing with the same concentrations of quinine (Seif et al., 2013, 2015). This may have allowed animals to desensitize to the same concentration of quinine over repeated exposures. Repeated tests with the same concentration were not employed in our study in order to 1) limit the possibility for quinine desensitization and 2) limit repeated injections of the chemogenetic actuator DCZ. Differences between our control group and those used previously may also warrant consideration. Whereas control subjects in Hopf et al. (2010) had continuous access to ethanol during the intermittent access period, our control subjects only had access to ethanol during a 5-day tapering protocol prior to testing. Low baseline levels of drinking in these controls may have limited our ability to observe suppression of drinking by quinine adulteration, preventing us from making a clear comparison to LTA rats. This may have also contributed to our inability to detect an aversion-resistant phenotype.

### Opposing effects of BF-to-LHb activation between sexes

In this study, we saw opposing effects of BF-to-LHb activation in male and female LTA rats. BF-to-LHb pathway activation reduced ethanol preference in males and promoted ethanol consumption in females. Given that stimulation of BF-to-LHb generally drives aversion (Faget et al., 2018; Swanson et al., 2022), while VP-to-LHb activation reduces cocaine seeking during cue-induced reinstatement in mice (Heinsbroek et al., 2020) and heroin seeking during relapse in male rats (Chen et al., 2024), we hypothesized that activation of this circuit would decrease ethanol intake-related behaviors. The effects of activation diminishing ethanol preference in males aligned with our hypothesis. We were surprised to find that activation of pathway promoted ethanol consumption in females. Opposing effects on behavior could be explained by distinct cell type profiles in this pathway. The majority of BF-to-LHb projections are glutamatergic (Barker et al., 2017; Faget et al., 2018; Levi et al., 2020). Glutamatergic projections from the BF to the LHb (Swanson et al., 2022) and VP to the LHb (Faget et al., 2018) promote place avoidance in mice, and inhibition of glutamatergic BF-to-LHb projections reduces social fear responses in mice (Wang et al., 2024). While many studies have focused on the role of glutamatergic BF output to the LHb, GABAergic and glutamatergic LPO-to-LHb projections both encode noxious stimuli (Barker et al., 2017), and parvalbumin-expressing VP-to-LHb neurons co-expressing both GABA and glutamate regulate depression-like phenotypes (Knowland et al., 2017), illustrating that overlapping cell type populations may be important in a variety of behaviors. While Barker et al. (2017) found that the majority (∼79%) of LPO neurons projecting to the LHb in male Sprague Dawley rats are glutamatergic, with a minority (∼4%) co-expressing GABA markers and a much smaller (∼16%) exclusively GABAergic population, Xu et al. (2022) found that LPO-to-LHb projections in male and female mice are more mixed. While ∼73% are glutamatergic, almost half of these (∼32% overall) also express GABAergic markers, and the remaining neurons appear to be GABAergic but not glutamatergic. Variability of the ratio of excitatory and inhibitory neurons in this pathway by species, strain, and sex, could lead to differences in the behavioral effects caused by activation. Thus, the effects observed here in females could be explained by a greater proportion of BF-to-LHb GABAergic neurons, or greater GABA release in the LHb, leading to additional ethanol consumption. Indeed, many previous studies examining the contributions of this pathway used only male subjects (Chen et al., 2024; Cui et al., 2022; Golden et al., 2016; Wang et al., 2024), including in observed reductions of heroin seeking during relapse after VP-to-LHb stimulation (Chen et al., 2024). This leaves investigation of this pathway in females relatively understudied. Future studies using cell-type specific approaches to characterize the neurochemical identity of these neurons in females versus males could help to shed more light on the sex differences reported here. Nevertheless, the sex-biased findings observed in the current study remain relatively modest, and more studies examining the effect of BF-to-LHb activation in both sexes would be useful to clarify the role of this pathway in the context of ethanol-related behaviors.

### BF-to-LHb activation suppresses consumption of quinine-adulterated sucrose

We also found that BF-to-LHb activation reduced suppression of sucrose intake by quinine. This aligned with findings from ethanol testing in females, where we observed an increase in ethanol intake following BF-to-LHb stimulation, but opposed findings in males where activation decreased ethanol preference. These results could be explained by differential recruitment of neurons in this pathway during consumption of sucrose when compared to ethanol. Selective pharmacological inhibition of the posterior VP, which comprises a higher relative proportion of GABAergic neurons versus its anterior subdivision (Faget et al., 2018), has been shown to eliminate “liking” reactions towards oral sucrose infusions, strongly inducing Fos expression in the LHb after sucrose exposure when compared to vehicle (>1100%) (Khan et al., 2020). This supports the idea that sucrose may preferentially engage GABAergic VP-to-LHb signaling to silence aversive signaling within the LHb. Therefore, chemogenetic activation of this pathway may further enhance LHb suppression, attenuating the effects of quinine on reducing sucrose intake observed in our experiments. Ethanol, however, may recruit distinct cell types within the BF. Prior research observed that chemogenetic inhibition of GABAergic VP neurons reduced context-induced reinstatement of alcohol seeking but did not reduce reacquisition of alcohol seeking upon reintroduction of alcoholic beer reward (Prasad et al., 2020). Meanwhile, inhibition of PV-expressing VP neurons (known to co-express glutamate and GABA and project to the LHb; Knowland et al., 2017) reduced both reinstatement and reacquisition of alcohol seeking (Prasad et al., 2020). These data suggest a potential shift towards glutamatergic VP signaling during ethanol versus sucrose consumption, implicating a potential mechanism for differences in downstream VP-to-LHb signaling. Recruitment of alternative ensembles varying in balance of excitatory and inhibitory signaling may have led to the differences we saw following BF-to-LHb chemogenetic activation during consumption of each reward. Although this mechanism is plausible, VP-to-LHb disconnection via contralateral lesioning did not affect either home-cage or operant ethanol intake (Sheth et al., 2017). However, due to its lack of cell-type specificity and ethanol-dependent signaling resolution, this lesioning approach may have obscured opposing GABAergic and glutamatergic contributions that carefully regulate ethanol drinking behavior. These neuronal populations could also respond differently to chemogenetic activation.

### Limited effects of BF-to-LHb inhibition

While chemogenetic excitation of BF-to-LHb projections yielded several interesting findings, inhibition did not elicit any observed effects on ethanol or sucrose drinking behaviors. This may not be entirely surprising when considering existing evidence of a more robust behavioral effect of BF-to-LHb activation compared to inhibition. While optogenetic stimulation of GABAergic BF-to-LHb inputs resulted in reduced capacity for impulse control, Hwang et al. (2024) found that BF-to-LHb optogenetic inhibition had no effect on this behavior, illustrating that inhibition of this circuit may not always yield an opposing response to activation. An important consideration remains that inhibition will only have an effect if endogenous activity of a target population is sufficient under behavioral circumstances; if BF-to-LHb neurons are relatively inactive during drinking tests, inhibition may not be expected to have any effect. Our results suggest that BF-to-LHb may not be necessary for normal alcohol and sucrose consumption or the effects of quinine. Other inputs to the LHb may play a greater role. For instance, entopeduncular nucleus projections to the LHb respond to reward- and aversion-predictive cues, pharmacological inhibition of which is sufficient to reinvigorate LHb signaling during cues predicting a sucrose reward (Li et al., 2019), while genetic inhibition can diminish stress-induced reinstatement to cocaine (Meye et al., 2016). Meanwhile, Sheth et al. (2017) found that lesioning lateral hypothalamus input to the LHb was capable of exacerbating home-cage ethanol intake.

## Conclusions

Here, we found that activation of BF-to-LHb circuitry diminished ethanol preference in male Long-Evans and elevated ethanol intake in female Long-Evans rats that had previously been exposed to long-term IAA, while reducing quinine-induced suppression of sucrose intake similarly between sexes. We found no significant effects of BF-to-LHb inhibition. To our knowledge, this is the first report of differential sex-specific effects of BF-to-LHb activation on consumption of alcohol or other drugs. An important next step will be to ascertain the roles of excitatory and inhibitory subpopulations within this pathway in these behaviors, as well as between distinct subregions within the BF. Future studies using a more robust aversion-resistant drinking model, incorporating genetic models sufficient to parse cell type and investigating subregion-specific BF-to-LHb projections, would help to disambiguate the effects of this pathway on ethanol drinking behaviors.

## SUPPLEMENTAL FIGURES

**Figure S1.**
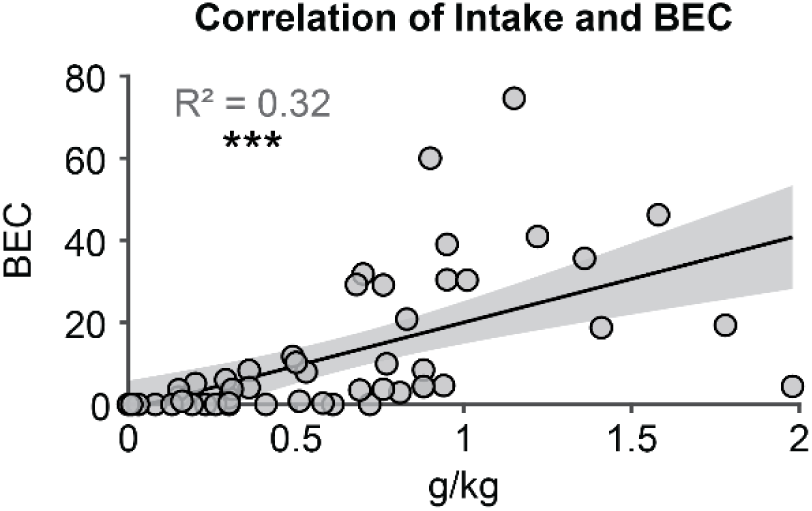
Blood ethanol content (BEC) correlates strongly with 30-minute g/kg ethanol intake. n = 50 total (25F/25M); 22 LTA (11F/11M) and 28 STA (14F/14M).

**Figure S2.**
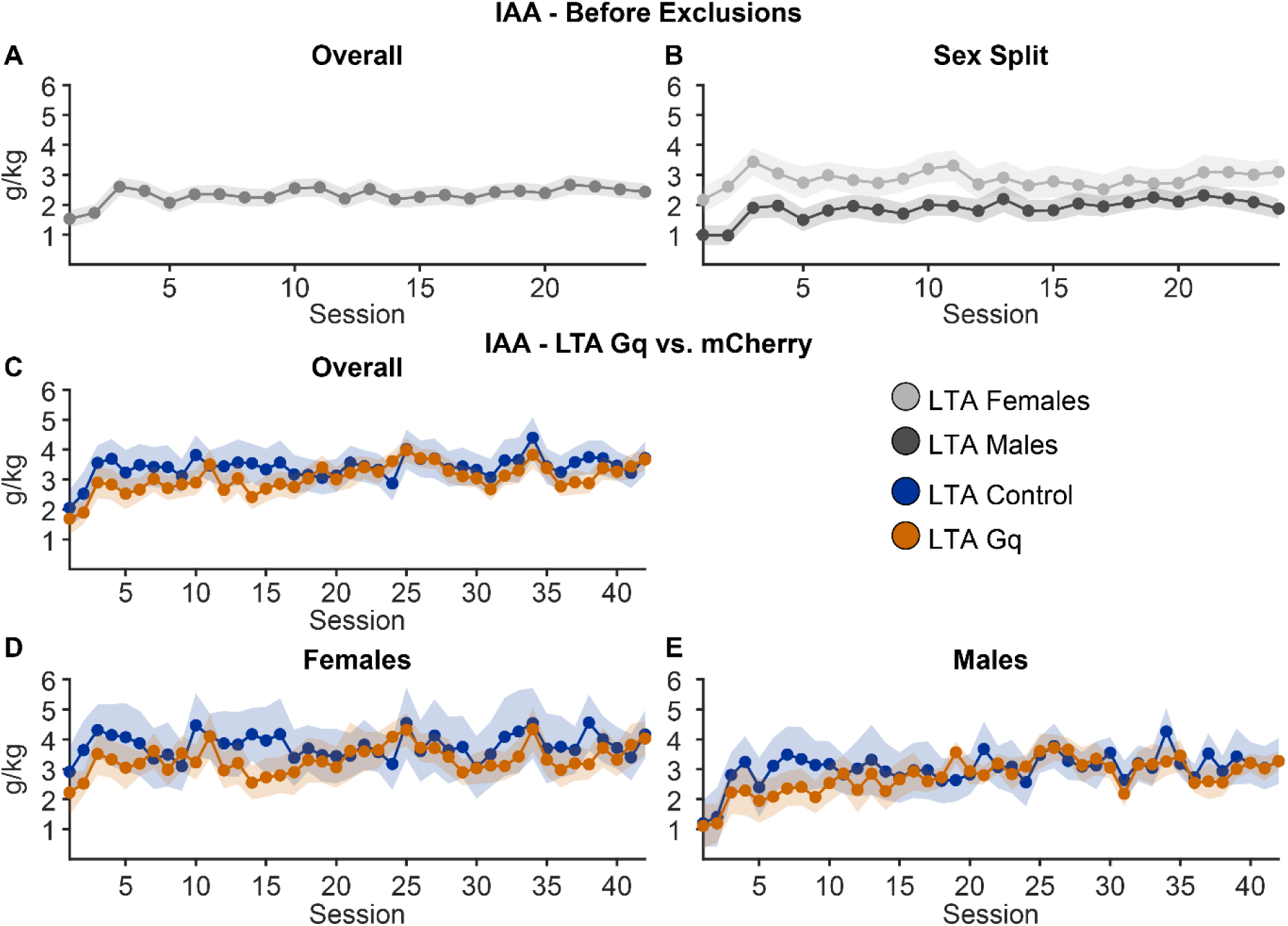
IAA analysis before and after intake exclusions. (A) IAA g/kg ethanol intake across all animals, pooling viral groups, prior to intake exclusions (IAA sessions 1-24) overall (n = 50), and (B) split by females (light grey; n = 23) and males (dark grey; n = 27). (D-E) IAA g/kg ethanol intake comparing between LTA control (blue; n = 14; 7F/7M) and LTA Gq (orange; n = 16; 8F/8M) animals after exclusions overall, and separately in (D) females and (E) males. *p< 0.05. Data are presented as mean ± SEM.

**Figure S3.**
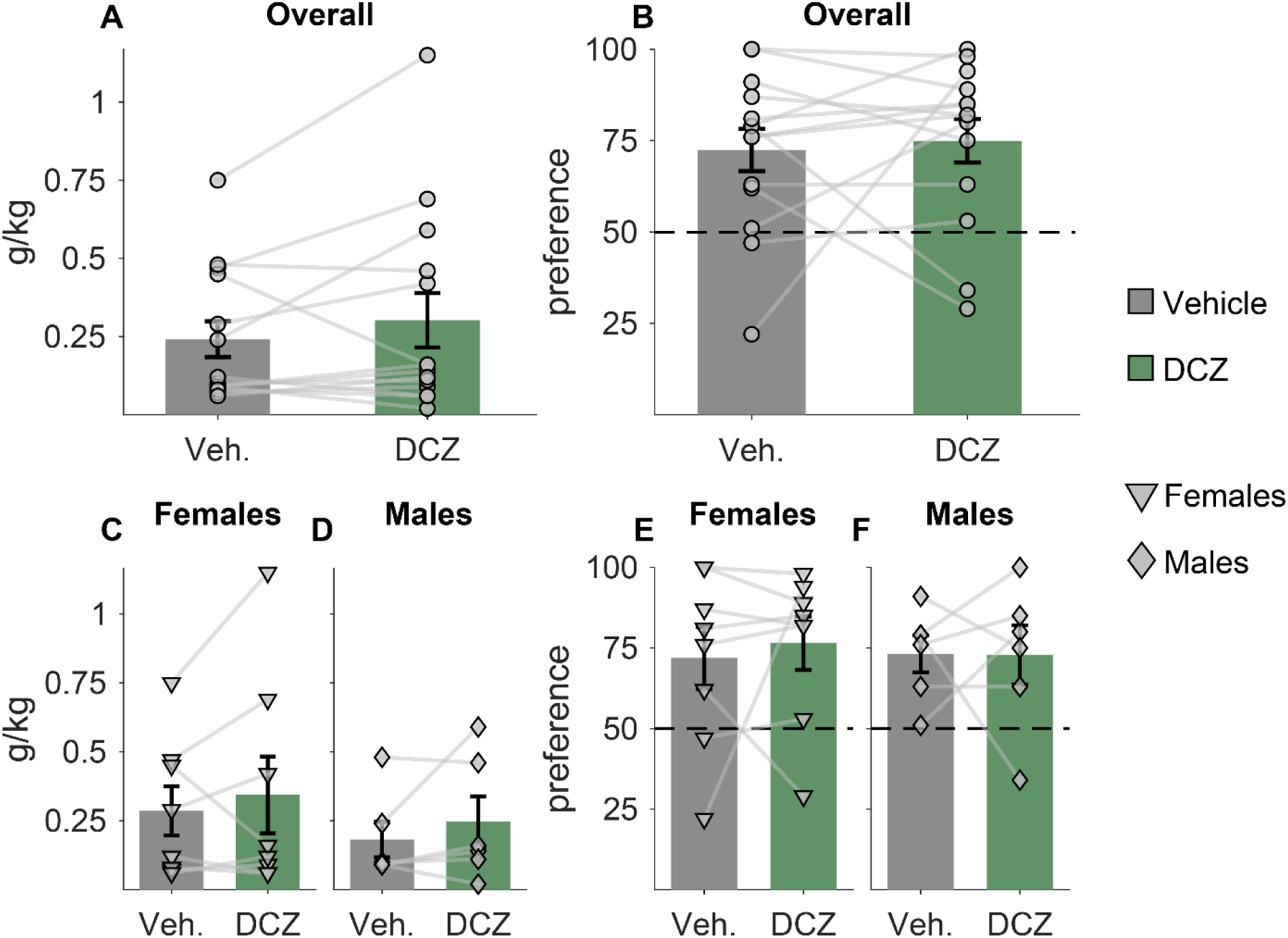
DCZ does not impact 30-minute ethanol intake in the absence of designer receptor expression. (A) Ethanol intake (g/kg) and (B) ethanol preference following saline (grey) and DCZ (green) injections in animals that did not receive virus (n = 14). Ethanol intake (g/kg) intake plotted separately for females (C; n = 8) and males (D; n = 6). Ethanol preference plotted separately for females (E) and males (F).

**Figure S4.**
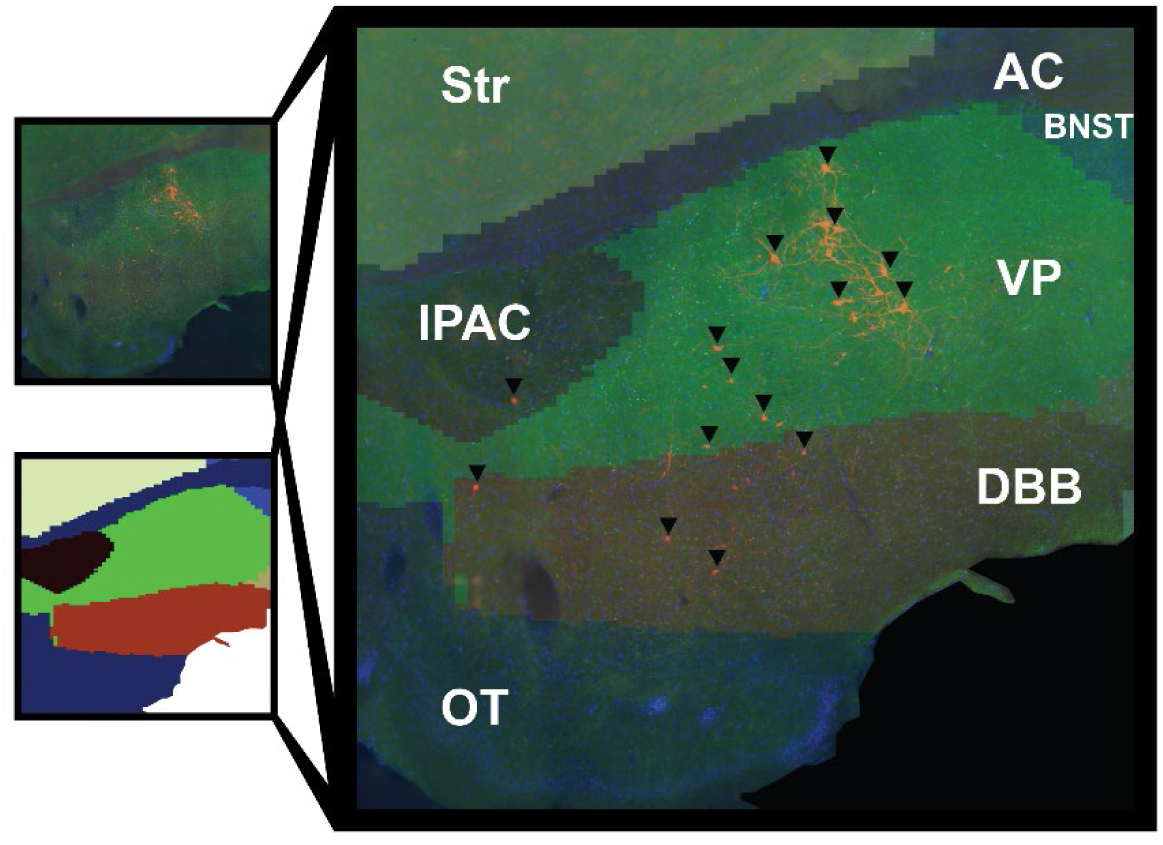
Example QUINT atlas alignment workflow image. VP = ventral pallidum; DBB = diagonal band of Broca; OT = olfactory tubercles; BNST = basal nucleus of the stria terminalis; Str = striatum; IPAC = interstitial nucleus of the posterior limb of the anterior commissure; AC = anterior commissure. Protocol based on *Yates, 2019*.

**Figure S5.**
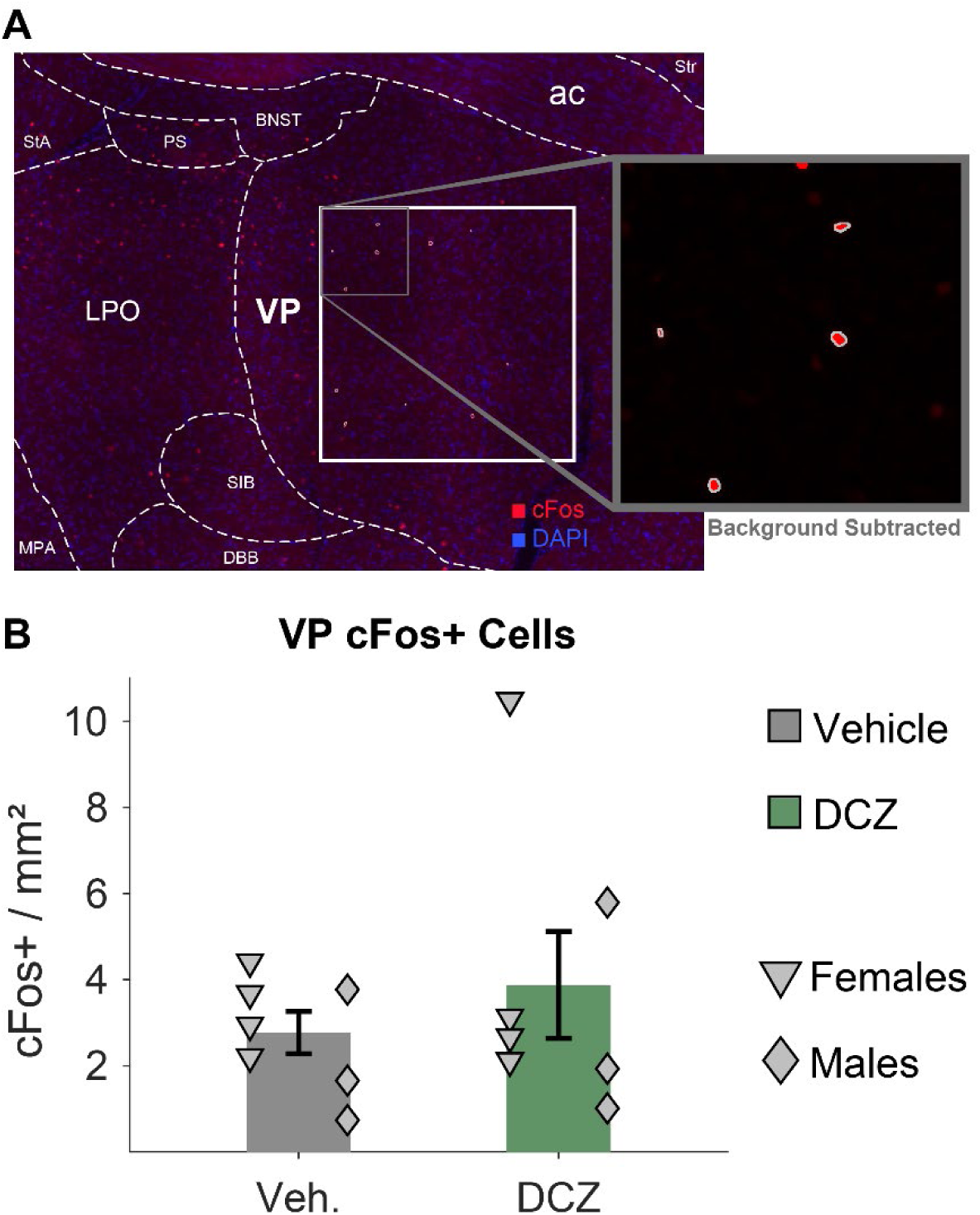
DCZ does not impact VP cFos expression in the absence of designer receptor expression. (A) Representative image of the basal forebrain with overlayed boundary designated for VP cFos quantification and (B) quantification of cFos level in ROIs drawn through the VP boundaries for saline vehicle (triangles, females, n = 4; diamonds, males, n = 3) and DCZ (females, n = 4; males, n = 3). VP = ventral pallidum; DBB = diagonal band of Broca; SIB = substantia innominata; BNST = basal nucleus of the stria terminalis; Str = striatum; LPO = lateral preoptic nucleus; MPA = medial preoptic area; PS = parastrial nucleus; StA = strial part of preoptic area; ac = anterior commissure.

## REFERENCES

Angelyn, H., Loney, G. C., & Meyer, P. J. (2021). Nicotine Enhances Goal-Tracking in Ethanol and Food Pavlovian Conditioned Approach Paradigms. Frontiers in Neuroscience, 15, 561766. 10.3389/fnins.2021.561766

Arnold, M. E., Decker Ramirez, E. B., Beugelsdyk, L. A., Siano Kuzolitz, M. V., Jiang, Q., & Schank, J. R. (2023). Estradiol mediates sex differences in aversion-resistant alcohol intake. Frontiers in Neuroscience, 17, 1282230. 10.3389/fnins.2023.1282230

Barker, D. J., Miranda-Barrientos, J., Zhang, S., Root, D. H., Wang, H.-L., Liu, B., Calipari, E. S., & Morales, M. (2017). Lateral Preoptic Control of the Lateral Habenula through Convergent Glutamate and GABA Transmission. Cell Reports, 21(7), 1757–1769. 10.1016/j.celrep.2017.10.066

Burchi, E., Makris, N., Lee, M. R., Pallanti, S., & Hollander, E. (2019). Compulsivity in Alcohol Use Disorder and Obsessive Compulsive Disorder: Implications for Neuromodulation. Frontiers in Behavioral Neuroscience, 13, 70. 10.3389/fnbeh.2019.00070

Calabrese, E., Badea, A., Watson, C., & Johnson, G. A. (2013). A quantitative magnetic resonance histology atlas of postnatal rat brain development with regional estimates of growth and variability. NeuroImage, 71, 196–206. 10.1016/j.neuroimage.2013.01.017

Carnicella, S., Ron, D., & Barak, S. (2014). Intermittent ethanol access schedule in rats as a preclinical model of alcohol abuse. Alcohol, 48(3), 243–252. 10.1016/j.alcohol.2014.01.006

Carpio, M. J., Gao, R., Wooner, E., Cayton, C. A., & Richard, J. M. (2022). Alcohol availability during withdrawal gates the impact of alcohol vapor exposure on responses to alcohol cues. Psychopharmacology, 239(10), 3103–3116. 10.1007/s00213-022-06192-8

Chen, H., & Lasek, A. W. (2020). Perineuronal nets in the insula regulate aversion-resistant alcohol drinking. Addiction Biology, 25(6), e12821. 10.1111/adb.12821

Chen, R., Liu, J., Wang, Y., Ning, K., Liu, J., & Liu, Z. (2024). Glutamatergic neurons in ventral pallidum modulate heroin addiction via epithalamic innervation in rats. Acta Pharmacologica Sinica. 10.1038/s41401-024-01229-4

Cofresí, R. U., Lee, H. J., Monfils, M.-H., Chaudhri, N., & Gonzales, R. A. (2018). Characterizing conditioned reactivity to sequential alcohol-predictive cues in well-trained rats. Alcohol, 69, 41–49. 10.1016/j.alcohol.2017.11.034

Cofresí, R. U., Lewis, S. M., Chaudhri, N., Lee, H. J., Monfils, M.-H., & Gonzales, R. A. (2017). Postretrieval Extinction Attenuates Alcohol Cue Reactivity in Rats. Alcoholism: Clinical and Experimental Research, 41(3), 608–617. 10.1111/acer.13323

Cofresí, R. U., Monfils, M.-H., Chaudhri, N., Gonzales, R. A., & Lee, H. J. (2019). Cue-alcohol associative learning in female rats. Alcohol, 81, 1–9. 10.1016/j.alcohol.2019.03.003

Cui, Y., Huang, X., Huang, P., Huang, L., Feng, Z., Xiang, X., Chen, X., Li, A., Ren, C., & Li, H. (2022). Reward ameliorates depressive-like behaviors via inhibition of the substantia innominata to the lateral habenula projection. Science Advances, 8(27), eabn0193. 10.1126/sciadv.abn0193

Faget, L., Zell, V., Souter, E., McPherson, A., Ressler, R., Gutierrez-Reed, N., Yoo, J. H., Dulcis, D., & Hnasko, T. S. (2018). Opponent control of behavioral reinforcement by inhibitory and excitatory projections from the ventral pallidum. Nature Communications, 9(1), 849. 10.1038/s41467-018-03125-y

Golden, S. A., Heshmati, M., Flanigan, M., Christoffel, D. J., Guise, K., Pfau, M. L., Aleyasin, H., Menard, C., Zhang, H., Hodes, G. E., Bregman, D., Khibnik, L., Tai, J., Rebusi, N., Krawitz, B., Chaudhury, D., Walsh, J. J., Han, M.-H., Shapiro, M. L., & Russo, S. J. (2016). Basal forebrain projections to the lateral habenula modulate aggression reward. Nature, 534(7609), 688–692. 10.1038/nature18601

Groeneboom, N. E., Yates, S. C., Puchades, M. A., & Bjaalie, J. G. (2020). Nutil: A Pre- and Post-processing Toolbox for Histological Rodent Brain Section Images. Frontiers in Neuroinformatics, 14, 37. 10.3389/fninf.2020.00037

Groos, D., & Helmchen, F. (2024). The lateral habenula: A hub for value-guided behavior. Cell Reports, 43(4), 113968. 10.1016/j.celrep.2024.113968

Heimer, L., Alheid, G. F., & Zahm, D. S. (1993). Basal Forebrain Organization: An Anatomical Framework for Motor Aspects of Drive and Motivation. In Limbic Motor Circuits and Neuropsychiatry. CRC Press.

Heinsbroek, J. A., Bobadilla, A.-C., Dereschewitz, E., Assali, A., Chalhoub, R. M., Cowan, C. W., & Kalivas, P. W. (2020). Opposing Regulation of Cocaine Seeking by Glutamate and GABA Neurons in the Ventral Pallidum. Cell Reports, 30(6), 2018–2027.e3. 10.1016/j.celrep.2020.01.023

Hikosaka, O., Sesack, S. R., Lecourtier, L., & Shepard, P. D. (2008). Habenula: Crossroad between the Basal Ganglia and the Limbic System. The Journal of Neuroscience, 28(46), 11825–11829. 10.1523/JNEUROSCI.3463-08.2008

Hopf, F. W., Chang, S., Sparta, D. R., Bowers, M. S., & Bonci, A. (2010). Motivation for Alcohol Becomes Resistant to Quinine Adulteration After 3 to 4 Months of Intermittent Alcohol Self-Administration. Alcoholism: Clinical and Experimental Research, 34(9), 1565–1573. 10.1111/j.1530-0277.2010.01241.x

Hopf, F. W., & Lesscher, H. M. B. (2014). Rodent models for compulsive alcohol intake. Alcohol, 48(3), 253–264. 10.1016/j.alcohol.2014.03.001

Hu, H., Cui, Y., & Yang, Y. (2020). Circuits and functions of the lateral habenula in health and in disease. Nature Reviews Neuroscience, 21(5), 277–295. 10.1038/s41583-020-0292-4

Hwang, E.-K., Zapata, A., Hu, V., Hoffman, A. F., Wang, H.-L., Liu, B., Morales, M., & Lupica, C. R. (2024). Basal forebrain-lateral habenula inputs and control of impulsive behavior. Neuropsychopharmacology, 49(13), 2060–2068. 10.1038/s41386-024-01963-7

Khan, H. A., Urstadt, K. R., Mostovoi, N. A., & Berridge, K. C. (2020). Mapping excessive “disgust” in the brain: Ventral pallidum inactivation recruits distributed circuitry to make sweetness “disgusting.” *Cognitive, Affective*, & Behavioral Neuroscience, 20(1), 141–159. 10.3758/s13415-019-00758-4

Knowland, D., Lilascharoen, V., Pacia, C. P., Shin, S., Wang, E. H.-J., & Lim, B. K. (2017). Distinct Ventral Pallidal Neural Populations Mediate Separate Symptoms of Depression. Cell, 170(2), 284–297.e18. 10.1016/j.cell.2017.06.015

LeCocq, M. R., Sun, S., & Chaudhri, N. (2022). The role of context conditioning in the reinstatement of responding to an alcohol-predictive conditioned stimulus. Behavioural Brain Research, 423, 113686. 10.1016/j.bbr.2021.113686

Levi, L. A., Inbar, K., Nachshon, N., Bernat, N., Gatterer, A., Inbar, D., & Kupchik, Y. M. (2020). Projection-Specific Potentiation of Ventral Pallidal Glutamatergic Outputs after Abstinence from Cocaine. The Journal of Neuroscience, 40(6), 1276–1285. 10.1523/JNEUROSCI.0929-19.2019

Levi, L. A., Inbar, K., Tseiger, E., & Kupchik, Y. M. (2025). A ventral pallidal glutamatergic aversive network encodes abstinence from and reexposure to cocaine. Science Advances.

Li, H., Pullmann, D., & Jhou, T. C. (2019). Valence-encoding in the lateral habenula arises from the entopeduncular region. eLife, 8, e41223. 10.7554/eLife.41223

Lüscher, C., Robbins, T. W., & Everitt, B. J. (2020). The transition to compulsion in addiction. Nature Reviews Neuroscience, 21(5), 247–263. 10.1038/s41583-020-0289-z

Meye, F. J., Soiza-Reilly, M., Smit, T., Diana, M. A., Schwarz, M. K., & Mameli, M. (2016). Shifted pallidal co-release of GABA and glutamate in habenula drives cocaine withdrawal and relapse. Nature Neuroscience, 19(8), 1019–1024. 10.1038/nn.4334

Millan, E. Z., Kim, H. A., & Janak, P. H. (2017). Optogenetic activation of amygdala projections to nucleus accumbens can arrest conditioned and unconditioned alcohol consummatory behavior. Neuroscience, 360, 106–117. 10.1016/j.neuroscience.2017.07.044

Millan, E. Z., Reese, R. M., Grossman, C. D., Chaudhri, N., & Janak, P. H. (2015). Nucleus Accumbens and Posterior Amygdala Mediate Cue-Triggered Alcohol Seeking and Suppress Behavior During the Omission of Alcohol-Predictive Cues. Neuropsychopharmacology, 40(11), 2555–2565. 10.1038/npp.2015.102

Napier, T. C., Kalivas, P. W., & Hanin, I. (Eds.). (1991). The Basal Forebrain: Anatomy to Function (Vol. 295). Springer US. 10.1007/978-1-4757-0145-6

Pan, G., Zhao, B., Zhang, M., Guo, Y., Yan, Y., Dai, D., Zhang, X., Yang, H., Ni, J., Huang, Z., Li, X., & Duan, S. (2024). Nucleus Accumbens Corticotropin-Releasing Hormone Neurons Projecting to the Bed Nucleus of the Stria Terminalis Promote Wakefulness and Positive Affective State. Neuroscience Bulletin, 40(11), 1602–1620. 10.1007/s12264-024-01233-y

Post, R. J., Bulkin, D. A., Ebitz, R. B., Lee, V., Han, K., & Warden, M. R. (2022). Tonic activity in lateral habenula neurons acts as a neutral valence brake on reward-seeking behavior. Current Biology, 32(20), 4325–4336.e5. 10.1016/j.cub.2022.08.016

Prasad, A. A., Xie, C., Chaichim, C., Nguyen, J. H., McClusky, H. E., Killcross, S., Power, J. M., & McNally, G. P. (2020). Complementary Roles for Ventral Pallidum Cell Types and Their Projections in Relapse. The Journal of Neuroscience, 40(4), 880–893. 10.1523/JNEUROSCI.0262-19.2019

Puchades, M. A., Csucs, G., Ledergerber, D., Leergaard, T. B., & Bjaalie, J. G. (2019). Spatial registration of serial microscopic brain images to three-dimensional reference atlases with the QuickNII tool. PLOS ONE, 14(5), e0216796. 10.1371/journal.pone.0216796

Rieck, R. W., Nabors, C. C., Updyke, B. V., & Kratz, K. E. (1995). Organization of the basal forebrain in the cat: Localization of l-enkephalin, substance P, and choline acetyltransferase immunoreactivity. Brain Research, 672(1–2), 237–250. 10.1016/0006-8993(94)01367-Q

Rodd, Z. A., Bell, R. L., McQueen, V. K., Davids, M. R., Hsu, C. C., Murphy, J. M., Li, T.-K., Lumeng, L., & McBride, W. J. (2005). Prolonged Increase in the Sensitivity of the Posterior Ventral Tegmental Area to the Reinforcing Effects of Ethanol following Repeated Exposure to Cycles of Ethanol Access and Deprivation. The Journal of Pharmacology and Experimental Therapeutics, 315(2), 648–657. 10.1124/jpet.105.084350

Segal, D., Valyear, M. D., & Chaudhri, N. (2022). The role of context on responding to an alcohol-predictive cue in female and male rats. Alcohol, 99, 70–81. 10.1016/j.alcohol.2021.10.004

Seif, T., Chang, S.-J., Simms, J. A., Gibb, S. L., Dadgar, J., Chen, B. T., Harvey, B. K., Ron, D., Messing, R. O., Bonci, A., & Hopf, F. W. (2013). Cortical activation of accumbens hyperpolarization-active NMDARs mediates aversion-resistant alcohol intake. Nature Neuroscience, 16(8), 1094–1100. 10.1038/nn.3445

Seif, T., Simms, J. A., Lei, K., Wegner, S., Bonci, A., Messing, R. O., & Hopf, F. W. (2015). D-Serine and D-Cycloserine Reduce Compulsive Alcohol Intake in Rats. Neuropsychopharmacology, 40(10), 2357–2367. 10.1038/npp.2015.84

Sheth, C., Furlong, T. M., Keefe, K. A., & Taha, S. A. (2017). The lateral hypothalamus to lateral habenula projection, but not the ventral pallidum to lateral habenula projection, regulates voluntary ethanol consumption. Behavioural Brain Research, 328, 195–208. 10.1016/j.bbr.2017.04.029

Siciliano, C. A., Noamany, H., Chang, C.-J., Brown, A. R., Chen, X., Leible, D., Lee, J. J., Wang, J., Vernon, A. N., Vander Weele, C. M., Kimchi, E. Y., Heiman, M., & Tye, K. M. (2019). A cortical-brainstem circuit predicts and governs compulsive alcohol drinking. Science, 366(6468), 1008–1012. 10.1126/science.aay1186

Simms, J. A., Steensland, P., Medina, B., Abernathy, K. E., Chandler, L. J., Wise, R., & Bartlett, S. E. (2008). Intermittent Access to 20% Ethanol Induces High Ethanol Consumption in Long– Evans and Wistar Rats. Alcoholism: Clinical and Experimental Research, 32(10), 1816–1823. 10.1111/j.1530-0277.2008.00753.x

Smith, K. S., Tindell, A. J., Aldridge, J. W., & Berridge, K. C. (2009). Ventral pallidum roles in reward and motivation. Behavioural Brain Research, 196(2), 155–167. 10.1016/j.bbr.2008.09.038

Sneddon, E. A., Schuh, K. M., Frankel, J. W., & Radke, A. K. (2021). The contribution of medium spiny neuron subtypes in the nucleus accumbens core to compulsive-like ethanol drinking. Neuropharmacology, 187, 108497. 10.1016/j.neuropharm.2021.108497

Soares-Cunha, C., & Heinsbroek, J. A. (2023). Ventral pallidal regulation of motivated behaviors and reinforcement. Frontiers in Neural Circuits, 17, 1086053. 10.3389/fncir.2023.1086053

Sood, A., Hanak, F., Joseph, A., Carpio, M. J., Gao, R., Selimovic, A., Weir, S. D. C., Uittenbogaard, F., Augustin, D., & Richard, J. M. (2026). Chronic intermittent ethanol vapor exposure alters local c-Fos and functional connectivity patterns evoked by alcohol cues. *Alcohol*, Clinical and Experimental Research, 50(2), e70258. 10.1111/acer.70258

Sparks, L. M., Sciascia, J. M., Ayorech, Z., & Chaudhri, N. (2014). Vendor differences in alcohol consumption and the contribution of dopamine receptors to Pavlovian-conditioned alcohol-seeking in Long-Evans rats. Psychopharmacology, 231(4), 753–764. 10.1007/s00213-013-3292-5

Swanson, J. L., Ortiz-Guzman, J., Srivastava, S., Chin, P.-S., Dooling, S. W., Hanson Moss, E., Kochukov, M. Y., Hunt, P. J., Patel, J. M., Pekarek, B. T., Tong, Q., & Arenkiel, B. R. (2022). Activation of basal forebrain-to-lateral habenula circuitry drives reflexive aversion and suppresses feeding behavior. Scientific Reports, 12(1), 22044. 10.1038/s41598-022-26306-8

Wang, J., Yang, Q., Liu, X., Li, J., Wen, Y.-L., Hu, Y., Xu, T.-L., Duan, S., & Xu, H. (2024). The basal forebrain to lateral habenula circuitry mediates social behavioral maladaptation. Nature Communications, 15(1), 4013. 10.1038/s41467-024-48378-y

Wise, R. A. (1973). Voluntary ethanol intake in rats following exposure to ethanol on various schedules. Psychopharmacologia, 29(3), 203–210. 10.1007/BF00414034

Xu, J., Jo, A., DeVries, R. P., Deniz, S., Cherian, S., Sunmola, I., Song, X., Marshall, J. J., Gruner, K. A., Daigle, T. L., Contractor, A., Lerner, T. N., Zeng, H., & Zhu, Y. (2022). Intersectional mapping of multi-transmitter neurons and other cell types in the brain. Cell Reports, 40(1), 111036. 10.1016/j.celrep.2022.111036

Yates, S. C., Groeneboom, N. E., Coello, C., Lichtenthaler, S. F., Kuhn, P.-H., Demuth, H.-U., Hartlage-Rübsamen, M., Roßner, S., Leergaard, T., Kreshuk, A., Puchades, M. A., & Bjaalie, J. G. (2019). QUINT: Workflow for Quantification and Spatial Analysis of Features in Histological Images From Rodent Brain. Frontiers in Neuroinformatics, 13, 75. 10.3389/fninf.2019.00075

Zhang, J.-P., Xu, Q., Yuan, X.-S., Cherasse, Y., Schiffmann, S. N., Kerchove d’Exaerde, A. D., Qu, W.-M., Urade, Y., Lazarus, M., Huang, Z.-L., & Li, R.-X. (2013). Projections of nucleus accumbens adenosine A2A receptor neurons in the mouse brain and their implications in mediating sleep-wake regulation. Frontiers in Neuroanatomy, 7. 10.3389/fnana.2013.00043

